# Nanopore Assay for Fingerprinting DNA Binding and Quantifying Real-Time Cleavage by Catalytically Active Cas9 Enzyme

**DOI:** 10.1101/2025.05.30.656938

**Authors:** Punitkumar Nagpure, Sarangi Suresh, Divya Shet, Gautam V. Soni

## Abstract

Nanopore sensing, a high-resolution DNA sequencing technology, is fast expanding into novel and exciting direction of probing specific DNA-enzyme interactions. Although proven excellent for detection of structural features of bare DNA, quantitative measurements on enzyme-DNA complexes and its real-time activity are lagging and only starting to emerge for long DNA templates. Signal-to-noise requirement and high translocation speeds make it difficult to detect protein bound on biologically relevant plasmid length DNA. To this end we report accurate position detection of a catalytically active Cas9 bound to its single or multiple target sites on the DNA. Protein position is fingerprinted using event charge deficit (ECD) based analysis of the high signal-to-noise electrical signals as the complex translocates through a glass nanopore. Using a time dependent assay, we quantify kinetics of the released products upon enzymatic cleavage of the target DNA by the wild-type Cas9 nuclease. Our approach enables the nanopore based single molecule sensing of DNA-protein complexes, for real-time monitoring of biochemical reactions. This may help understand protein binding & localization as well as improve Cas9 based targeting in genome engineering applications.

**TOC Figure:** 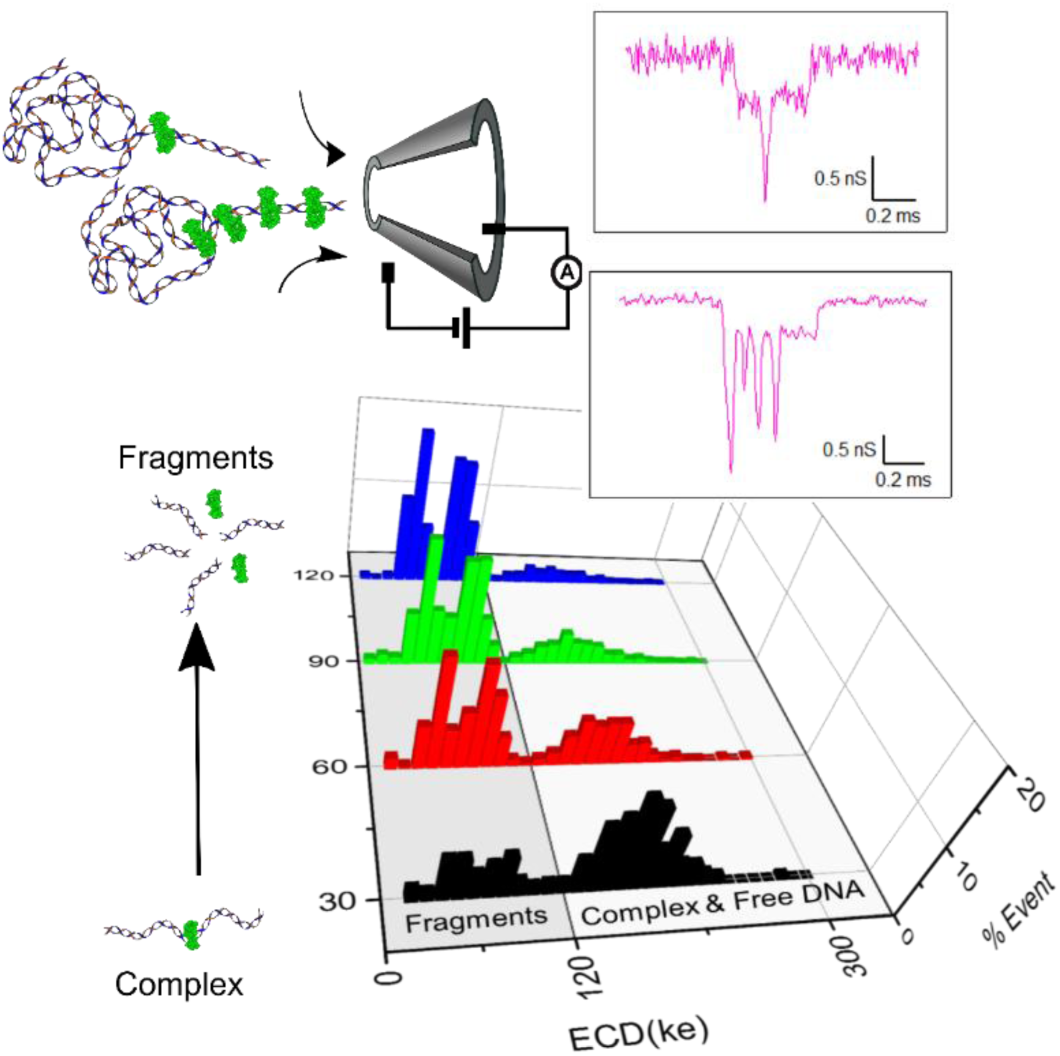

## INTRODUCTION

Solid state nanopore is an emerging single molecule measurement platform for probing DNA-protein complexes at molecular scale while providing high throughput detection. Nanopore sensors are high resolution, label-free detection technique that work on resistive pulse sensing. Nanopores are small (few nanometres) orifices in thin solid-state membranes or at the tip of glass nanopipettes^1–8^. The analyte molecule, in electrolyte buffer, electrophoretically translocates through a nanopore under an applied voltage. This causes temporary blockades in nanopore current for the translocation duration. These electrical events form typical nanopore signal, characterized by conductance blockade (ΔG) and translocation time (Δt) which correspond, respectively, to the dimensions and the charge or length of the analyte molecule^1^. Recently, nanopore platform has been utilized to detect DNA as well as other biological nanostructures^5,9–14^ There are also recent nanopore studies of DNA-protein complexes that are detected , such as EcoRI, RecA, RNAP, Nuclesomes etc^6,15–21^.

DNA-protein interactions play a central role in various cellular and biochemical processes. These processes often involve binding and unbinding of various proteins at specific locations on DNA. Information on their position on the DNA can provide valuable insights into their role in regulating and maintaining genomic functions^22,23^ . In recent years, CRISPR (Clustered Regularly Interspaced Short Palindromic Repeats) technology has revolutionized the field of genome editing due to its remarkable ability to target specific sequences in the genome and its applications in diagnostics^24–26^. The CRISPR-Cas9 system relies on the Cas9 protein, guided by specific RNA molecules, to bind and cleave target DNA at precise locations. Given its transformative applications in genome editing and diagnostics, there is a significant interest in enhancing the specificity and the efficiency of CRISPR technology. Therefore, it is crucial to develop simple, high-throughput, and sensitive assays to measure the binding of Cas9 and its localization on the target DNA. A real-time report on its cleaved DNA products will also help to better understand and optimize the effectiveness of CRISPR systems.

The Cas9 is the catalytically active form of the CRISPR associated protein. It is a bi-lobed protein of size 159 kDa (from manufacturer) and dimensions 16×21×9.1 nm^27,28^. Target DNA (protospacer) usually consists of 20 bp sequence adjacent to a protospacer adjacent motif (PAM sequence, NGG). Cas9 enzyme is loaded with a duplex guide RNA (crRNA and tracrRNA) which complements and identifies the 20 bp target sequence on the template DNA. Efficient Cas9 binding to target DNA and its cleavage require canonical base paring between crRNA and target DNA sequences. The Cas9 can be targeted to a different location by simply changing the crRNA sequence. It has been shown that Cas9 remains stably bound to the target DNA after cleavage^29^ holding on to the cleaved products. The products can then be released with surfactant or heat^30–33^. Recent work has studied different aspects of CRISPR-Cas9 system such as binding ^33–35^, target search dynamics^36,37^,stability and off target activity^38–40^ by employing various single molecule tools such as AFM, optical tweezers, magnetic tweezer and florescence based imaging. Even though these techniques are excellent to study functional aspects of the CRISPR enzyme, quantifying precise physical properties of the DNA-enzyme system remains challenging due to the experimental complexity, chemical treatment of the sample and the required sensitivity.

Recent studies have addressed nanopore-based detection of the binding of various proteins including ZEP^41^, DNA–antibody complexes^19^, RNA polymerase^42^, and catalytically inactive Cas9^43,44^. These studies typically employed long DNA constructs, such as 20 kb – 48 kb linear DNA as substrates for protein binding. However, these long DNA molecules have a higher tendency to undergo folded translocations through the nanopore, which introduces complexity in localizing bound protein. Additionally, nanopore-based approaches have been used to monitor the cleavage products of Cas12a enzymes, utilizing DNA tetrahedron origami structures or circular single-stranded DNA as carrier molecules for the cleaved fragments^45–47^. More recent work has also demonstrated that polymer-based crowding agents can enhance nanopore sensitivity, enabling the monitoring of cleavage activity at low ionic strength conditions of 0.1 M KCl^48^. However, operating at lower salt concentrations presents several challenges, including non-linear current–voltage relationships, increased interactions between proteins and nanopore walls, and reduced resolution for accurately detecting and characterizing DNA–protein complexes such as binding position, protein volume and enzymatic products.

In this work, we first establish a complete work plan to accurately quantify different DNA lengths from a mixture using our conical glass nanopore system and a detailed event charge deficit (ECD) based analysis. This allowed us to identify the lengths of the translocating DNA molecule with 189 bp resolution (in < 2 kbp range) . We then demonstrate identification of Cas9 complexed DNA molecules translocating through the nanopore, where the Cas9 bound DNA complexes were distinctly identified by the presence of secondary current spikes in the nanopore signal. We employ ECD analysis to localize position of the a centrally-located Cas9 enzyme on the DNA with better than 4 % error. Owing to our high signal-to-noise, we further demonstrate detection of multiple Cas9 molecules bound to a multi-target DNA. Finally, we apply the ECD-based DNA length analysis to measure real-time product release after cleavage of the target DNA by the catalytically active Cas9 enzyme.

This work expands the high-resolution detection of nanopore platform from DNA sequencing applications to single molecule enzymatic assays. We envision this as a first step in many new potential applications of measuring specific and non-specific protein binding and functional enzymatic assays, monitoring assembly and disassembly of multi-protein & DNA complexes.

## MATERIALS AND METHODS

### Nanopore Fabrication and Experiment

We used a laser assisted glass capillary puller (P-2000G, Sutter Instrument) to fabricate glass (quartz) nanopores, as described previously^4,5^. Briefly, the capillaries (0.65 mm (OD) x 0.35 mm (ID), Sutter Instrument) were pulled with following two-line program:

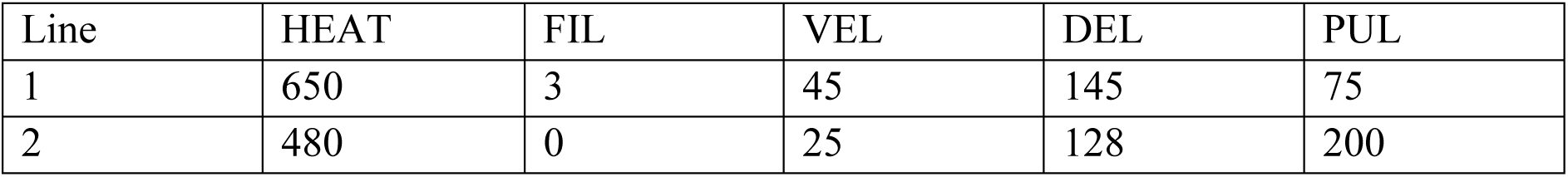

| Line | HEAT | FIL | VEL | DEL | PUL |
| --- | --- | --- | --- | --- | --- |
| 1 | 650 | 3 | 45 | 145 | 75 |
| 2 | 480 | 0 | 25 | 128 | 200 |

This capillary pulling program results in pulled capillaries with nanometer sized (50 – 80 nm) pores at its tip. We then shrink these nanopores to desired pore diameters (∼20 nm in this study) by irradiating the tips with an electron beam (3 keV) from the Scanning Electron Microscope (SEM, Carl Zeiss Ultraplus FESEM). Final nanopore diameters and taper dimensions of the capillary were quantified by SEM and optical imaging, respectively (see Figure 1A). Before each experiment these quartz nanopores were cleaned in oxygen plasma using Plasma Prep-III system (SPI Supplies) to make them hydrophilic and then mounted in a custom-made Teflon fluid cell using Ecoflex 005 glue (Smooth-On Inc.). The fluid cell has two chambers and the capillary with the nanopore was mounted across the two chambers (see schematic in Figure 1B). The chamber with nanopore mouth (*cis*-chamber) was filled with nanopore buffer (NPB) containing 2 M LiCl in 10 mM Tris-HCl, 1 mM EDTA with pH 8 and left for 5 min to let the buffer fill the nanopore and its conical taper by capillary action. Then NPB was added into the *trans* chamber filling the back side of the capillary. The fluid cell with the nanopipette was then placed in a desiccator for 10 min to remove any air bubbles. I-V curves and open pore current was recorded to check for linearity and stable low-noise (< 5 pA) baseline behavior before adding sample. Ag-AgCl electrodes and Axopatch 200B amplifier (Molecular Devices) were used for all current measurements in this work. Typical sample concentrations (to minimize pore blockage) used in translocation experiments are 0.8 nM for free DNA and 0.07 – 0.14 nM for Cas9-DNA complex. The hardware bandwidth on the amplifier was set to 100 kHz and the data was acquired at a sampling rate of 200 kHz using NI-PCI 6251 DAQ card. Custom written LabVIEW (National Instruments) codes were used for data acquisition and further analysis. Current data was low-pass filtered with 25 kHz cutoff for all event detection and analysis.

**Figure 1:**
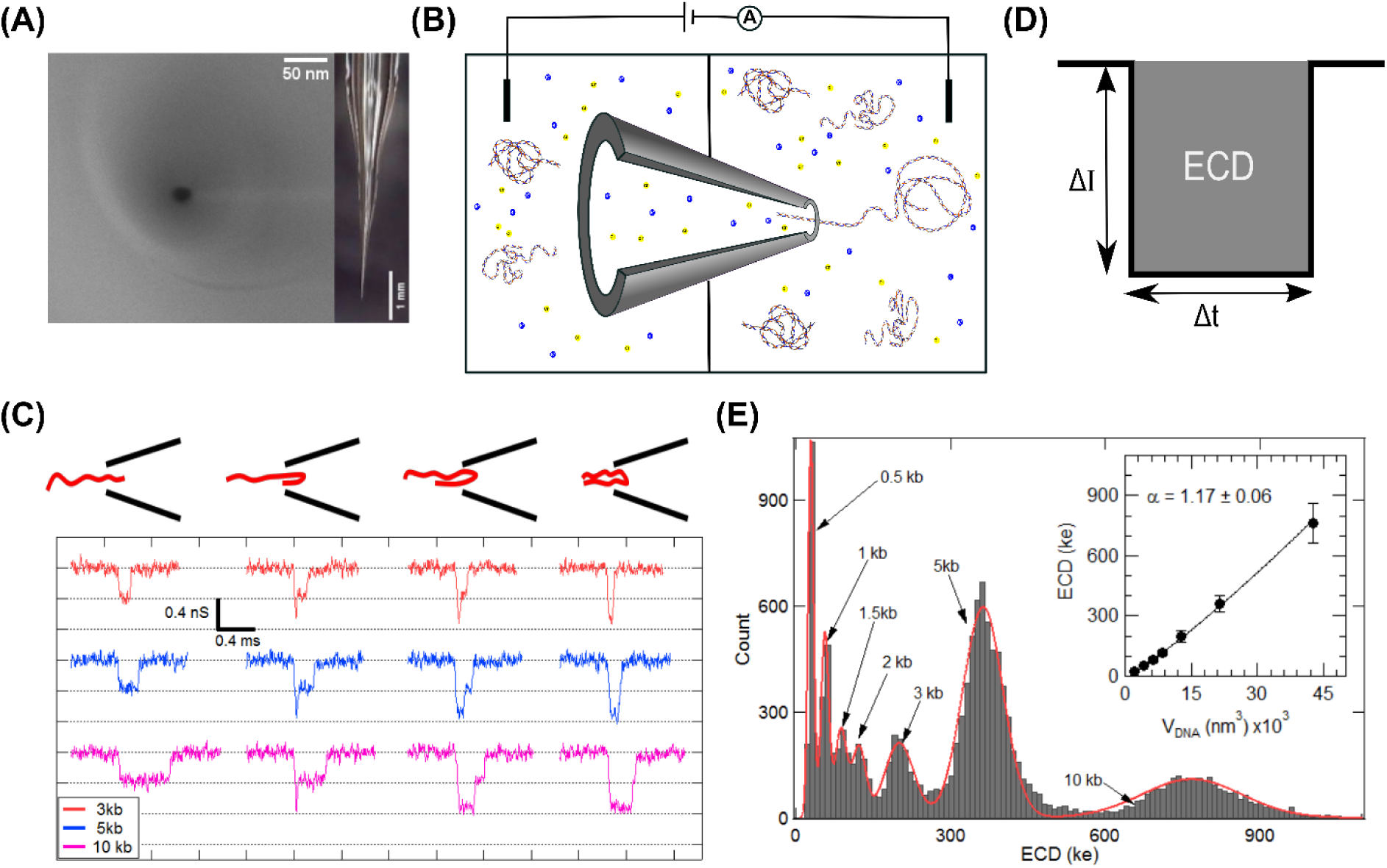
Detection and characterization of different sizes of linear DNA in glass nanopore. (A) Left: SEM image of a typical 20 nm diameter glass nanopore. Right: Optical side-on image of the tapered glass capillary. (B) Schematic of the experimental setup showing glass nanopore mounted on Teflon fluid cell along with buffer (yellow and blue ions) solution and sample (DNA). The voltage is applied through Ag-AgCl electrodes and current is measured using the amplifier. (C) From the custom-mix of DNA, representative events corresponding to 3,5, & 10 kb DNA translocating through the 20nm nanopore is shown. Different possible folded configurations corresponding to the events are shown at top. (D) Schematic showing typical translocation event parameters of current blockade (ΔI) and the dwell time of the event (Δt). The gray shaded area of the event measures the event charge deficit (ECD). (E) ECD histogram from events (N=14391) of custom-mix sample of DNA molecules (sizes range 0.5 – 10 kb) translocating through 20 nm nanopore. The inset shows the relationship between ECD peak values and volume (πrDNA^2^LDNA) of the translocating DNA molecule. The solid line is a power-law fit with exponent α = 1.17.

DNA sensing and sizing capabilities of our nanopores were tested using our custom-mix DNA sample. This was made by adding 5 and 10 kb linear plasmid DNA to a 1 kb DNA ladder (NEB N3232S). Final concentrations of each DNA length in the custom-mix are shown in Table S1.

### Preparation of Target DNA for Cas9 binding

The target DNA was pGEM-3z/601 plasmid (3025 bp) with the target sequence as: 5′-GGCACCGGGATTCTCCAGGG-3′. Two distinct position of the same target sequence was achieved by linearizing the plasmid DNA with two different single-cut restriction enzymes. The plasmid linearized with ScaI-HF (NEB, R3122S) enzyme resulted in Cas9 target sequence at 1853 bp from the cut end. This target DNA is referred to as “center-target” in the study. Linearization with the restriction enzyme EcoRI-HF (R3101S) resulted in Cas9 target to be at 215 bp from the cut site. This target is referred to as “edge target” in this study. For multiple Cas9 binding to DNA, we used a 5094 bp linear DNA (linearized from pUC18-12×601 plasmid, from Dekker lab) which has 12 repeats of the same target sequence on one half of the DNA (see Figure 4A).

### Duplex RNA Formation

The CRISPR RNA (crRNA) and *trans*-activating CRISPR RNA (tracrRNA), corresponding to the target sequence, were purchased from Integrated DNA Technology (IDT, USA). They were resuspended in nuclease free buffer (30 mM HEPES, 100 mM potassium acetate, pH 7.5, provided by manufacturer) to make 100 µM stock of each RNA. The formation of the duplex RNA was achieved by mixing crRNA and tracrRNA in nuclease-free buffer at an equimolar concentration, resulting in a final duplex guide RNA of 1 µM. The mixture was heated to 90 °C for 30 s, then slowly cooled to room temperature (1 h) to yield the crRNA-tracrRNA duplex (RNA).

### Cas9-RNA and DNA binding reaction

The Cas9 Nuclease, *S. pyogenes* for the experiments were purchased from New England Biolabs (NEB, M0386S). RNA loaded Cas9 were made by mixing the duplex RNA and Cas9 enzyme in 1:1 molar ratio at 60 nM concentration and incubated for 15 min at 37 °C in HEPES buffer (25 mM HEPES-NaOH (pH 8.0), 150 mM NaCl, 1 mM MgCl_2_). The binding to the target DNA was achieved by adding the target DNA to the RNA loaded Cas9, maintaining the molar ratio of 1:30:30 (DNA: Cas9: RNA) and incubating for 1 h at 37 °C. After the reaction, samples were used immediately. We have previously shown^33^ that this reaction results in stable Cas9-DNA complexes.

The stability of the Cas9-DNA complex in NPB (with 2M LiCl) was verified using electrophoretic shift assay shown in Figure S1. The Cas9-DNA complex was incubated in 2M (LiCl) salt and tested for complex stability, for up to 2 hours. We note, that in the duration of 2 hours, the complex band reduces in intensity, gradually (Figure S1 lanes 9 – 12). This slow release of products, allows for measurement of Cas9-DNA complexed structures within our typical experiment duration (15 – 45 min). Note that, the release of cleaved products is discussed later in the text (see Figure 6). The cleaved products could be completely released from the complex by heating the sample to 90 °C for 10 min (see Lane 13 in Figure S1).

For multi Cas9-DNA complex, 5094 bp (5 kb) DNA and RNA loaded Cas9 were mixed in 1:40:40 ratio and incubated at 37 °C for 1 hour and then at 4 °C overnight to maximize the binding. This DNA to Cas9 ratio resulted in sub-saturated number of Cas9 bound to the possible 12 target sites of the 5 kb DNA.

## RESULTS

### Quantitative Nanopore Analysis of DNA sizing

First, we demonstrate the sensitivity of our nanopore platform in quantifying linear DNA of different lengths. We used the custom-mix of linear DNA (see material and methods) in the size range 0.5 kbp to 10 kbp, with the final concentration of each size of DNA tabulated in Table S1. We used nanopores of size ∼20 nm diameter for this study (see Figure 1A). After adding the DNA mixture into the experiment chamber translocation events were recorded at 500 mV as shown in schematics of Figure 1B. Typical example of the baseline current before and after adding sample are shown in Figure S2. Voltage driven translocation events of individual DNA molecules traversing through the nanopore are detected and for every event, parameters such as the current drop (ΔI) and the duration of translocation time (dwell time, Δt) (see definitions as schematics in Figure 1D and Figure S2D) is recorded. A representative selection of the translocation events is shown in Figure 1C. We see the multi-level events as seen previously^5^. These individual multi-level events correspond to unfolded (linear) as well as partially and fully folded conformations of DNA molecules as they translocate through the nanopore. As the number of DNA folds increases (from left to right in Figure 1C), the conductance decreases in well-defined steps (dotted lines). We also see that the translocation times increase with DNA length.

A quantitative identification of the length of the translocating DNA molecules, irrespective of its conformation, can be done by evaluating its event charge deficit (ECD). The integral of the current drop (ΔI) over the dwell time (Δt) duration for an event is termed as its event charge deficit (see schematic in Figure 1D). The ECD of each DNA molecule translocating the nanopore is proportional to the volume of ions displaced and independent of the conformation (linear or folded) in which the molecule translocated. In Figure 1, we used custom-mix DNA sample of sizes 0.5 – 10 kb. Each of these DNA lengths translocate through the nanopore (in various conformations) producing events with ECD characteristics corresponding uniquely to their size. We collected more than 10000 events and evaluated their ECDs (see histogram in Figure 1E). It shows that the mixture of DNA lengths translocating through the nanopore result in well separated ECD peaks with the peak values proportional to the respective DNA lengths. The peaks corresponding to specific DNA lengths were identified by correlating relative concentrations of DNA lengths to ECD peak heights. We compared ECD histograms of sample containing DNA lengths (0.5 kb to 3 kb) of 1 kb DNA ladder (see Figure S3A & S3B) to the custom-mix DNA sample and found new peaks emerging corresponding to the externally added 5 kb and 10 kb DNA lengths (see Figure 1E and Figure S3C & S3D). Average ECD peak positions (from multiple nanopore and sample concentrations experiments) corresponding to specific DNA lengths were thus identified and plotted in Figure 1E (inset) (see also Figure S3E & S3F and Table S2). Note that the DNA lengths with concentrations lower than 0.11 nM result in very low event frequency and hence are not detected as distinct peaks. The ECD versus DNA volume plot was fitted with power law equation ECD (ke) = A × (V_DNA_)^α^, where V_DNA_ is the volume of the translocating DNA molecule in nm^3^ and A is the fitting parameter^14^. The fitting reveals power law dependence of the ECD values on the volume of the translocating DNA. This curve can also be used as calibration to identify lengths of unknown DNA fragments in the sample. We have reproduced these results with multiple nanopores (see Figure S3) and the corresponding ECD values are shown in Table S2. The mean power-law exponent from multiple experiments (see Table S3) was found to be α = 1.17. Our results show the sensitivity of our nanopore device and ECD analysis in detecting DNA molecules of length down to 0.5 kb. From the spread of ECD histograms, we could estimate a length resolution of ∼ 189 bp for DNA lengths < 2 kbp (see Table S2 & S3).

### Detection of DNA-Cas9 Complex

The Cas9 proteins were bound on the linear 3025 bp DNA to the center target position and the stability of the complex in the experimental buffer (NPB – 2M LiCl) was confirmed by electrophoretic mobility shift assay (see material and methods and Figure S1). The control DNA (Figure 2A) and the Cas9 bound DNA (Cas9-DNA complex, Figure 2D) were translocated through the same nanopore, back-to-back, at 300 mV. The events collected for both samples were analyzed and the ΔG and Δt of each event is plotted as scatter plots in Figure 2B and 2E, as indicated. Control DNA translocates through the nanopore in linear or folded configuration (see Figure 1C). Population of events corresponding to the two configurations are seen as two population scatters in Figure 2B. Representative events of these two types of translocations are shown in Figure 2C (a) & (b). The mean event depth of linear and folded DNA was measured to be ΔG_1_ = 0.44 ± 0.15 nS and ΔG_2_ = 0.78 ± 0.18 nS, respectively (see Figure S4). We note that DNA in folded conformation results in blockade levels close to twice that of the unfolded DNA. These two configurations cover most of the translocation events in the case of control DNA sample.

**Figure 2:**
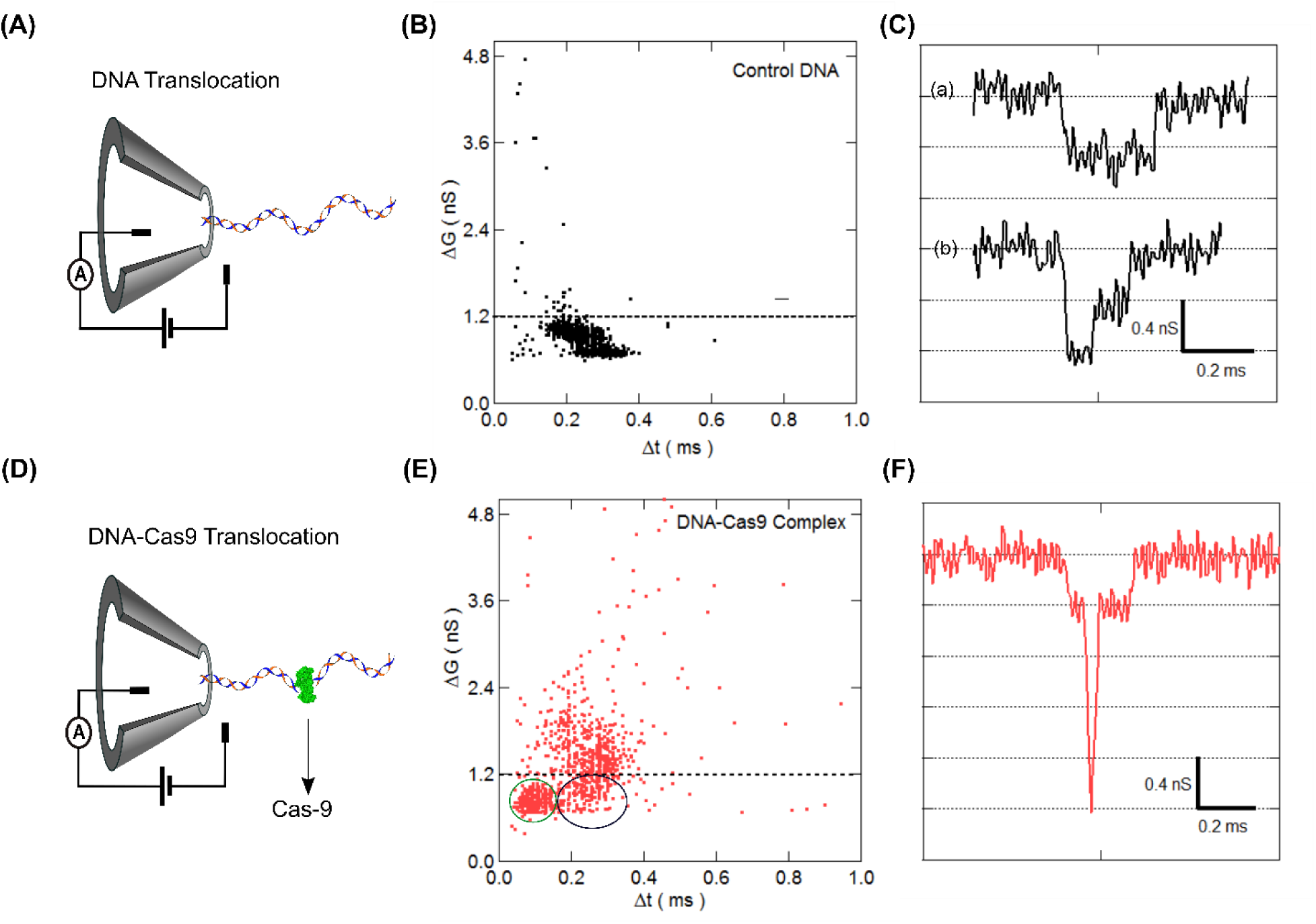
Detection of Cas9-DNA Complex. (A) and (D) shows schematics of translocation of control DNA and Cas9-DNA complex through nanopore, respectively. (B) and (E) shows scatter plot of conductance blockade (ΔG) verses dwell time (Δt) for control DNA (N = 1567 events) and Cas9-DNA complex sample (N = 589 events), respectively. Green and black circles in (E) represent cluster of events corresponding to cleaved fragments and unbound DNA respectively. The events above the dotted threshold line represent translocation of stably bound Cas9-DNA complex. (C) shows representative translocation event of the control DNA in linear (top, (a)) and partially folded (bottom, (b)) conformation. (F) shows representative event corresponding to translocation of a Cas9-DNA complex. The event shows current levels corresponding to the baseline, the DNA and the presence of the Cas9 protein bound on DNA, seen as a distinct spike on top of the DNA level.

In the same nanopore, translocation of Cas9-DNA complex was performed (see Figure 2D). The events collected were analyzed for ΔG and Δt values and are shown as scatter plot in Figure 2E. We note that other than the linear and folded population, the Cas9-DNA complex sample produced a large number of events with very high ΔG values. These events were considered for further analysis. We isolated these events using a simple threshold of ΔG > 3 × ΔG_1_ as shown in black dotted line in Figure 2E. This threshold of ΔG > 1.2 nS was chosen since, for control DNA only 0.05 % of the events were found to be above this threshold (see black dotted line in Figure 2B). These small number of events are primarily single-level events with very small dwell times which may be attributed to translocation of higher-level folding in DNA. Representative of such events is shown in Figure S5. Using this approach, we found that in a typical experiment, about 40 – 60 % of the events are above the threshold during the translocation of the Cas9-DNA complex. This was reproduced for multiple nanopores and sample preparations (see Figure S6 and Table S4). Closer inspection of these events shows a sharp and deep current spike on top of the linear DNA current blockade level, as shown in Figure 2F. We note that these spikes have short duration and large conductance drop. These events with the secondary spike were exclusively found during the measurement of Cas9-DNA complex sample. We attribute it to the presence of Cas9 protein bound on the translocating DNA. A library of such events is shown in Figures S8A & S8B. We observe the conductance drop due to the Cas9-DNA complex was ∼ 3.4 times the control DNA which is better than previous reports (see Figure S7)^44^. Additionally, we observe two distinct population below the threshold in scatter plot Figure 2E, visually marked with black and green circles. Upon closer inspection, the events marked with black circle were similar to control DNA events (Figure 2B), possibly representing the unbound population of control DNA. We also observe a distinct cluster of events with shorter dwell times shown in green circle. These events correspond to translocation of DNA shorter than the target control DNA and are possibly from the translocation of the released cleaved DNA fragments after the Cas9 enzymatic activity. Evaluation of these events by ECD analysis is shown later in the text.

### Localization of Cas9 bound on target DNA

Figure 3A (top-left) shows the schematic of the sample where the Cas9 is bound to the target site located towards the center of the linear DNA (see Figure 3A top-right & Materials and Methods section). This sample is referred to as center-target in the text. Analyzing many such events with the protein spike, we find that the all events have the spike located at two distinct locations along the event. Representative events of these two spike positions are shown in Figure 3A (right) and Figure S8B. We understand the two protein positions are result of the Cas9-DNA complex entering the nanopore in two different orientations (as shown in Figure 3A (left)). This results in the two possible protein positions measured on the translocating DNA.

**Figure 3:**
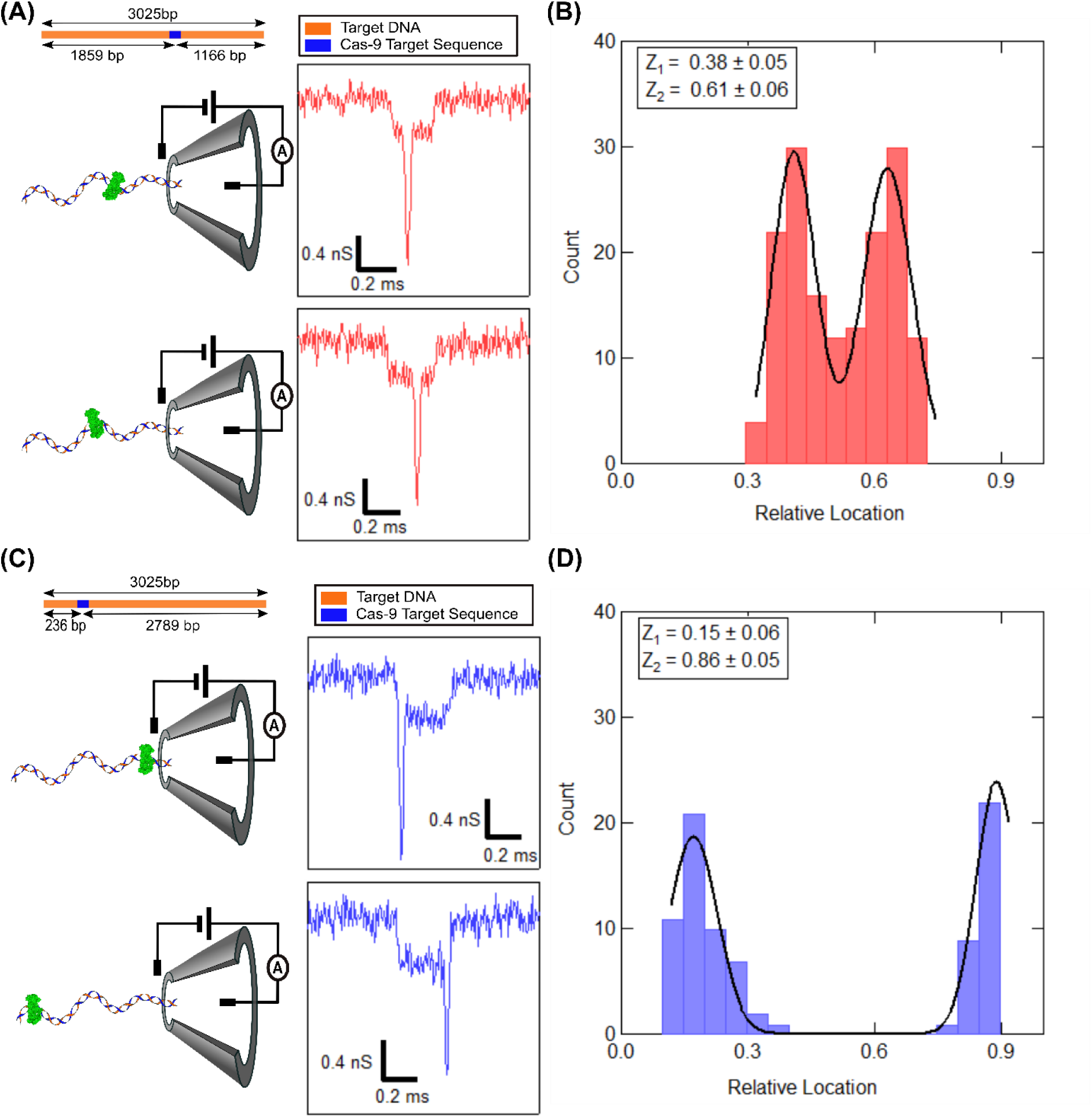
Spatial localization of Cas9 protein bound to the linear 3kb target DNA. (A) (top) shows the position of the target (blue) on the 3 kb DNA (orange), in the case of center-target. Lengths of the target DNA and the cleaved products are indicated. (Bottom) shows schematic (left) and representative event (right) depicting the two orientation in which the complex can translocate through the nanopore. (B) shows histogram of ECD-based estimation of the spike positions relative to the event length for the center-target sample. (C) and (D) are the same as above for DNA-Cas9 complex bound on the edge target. Z1 and Z2 are the peak positions that gives estimate of the relative position of the spike along the DNA event, where the DNA is translocating in the two different orientations.

**Figure 4:**
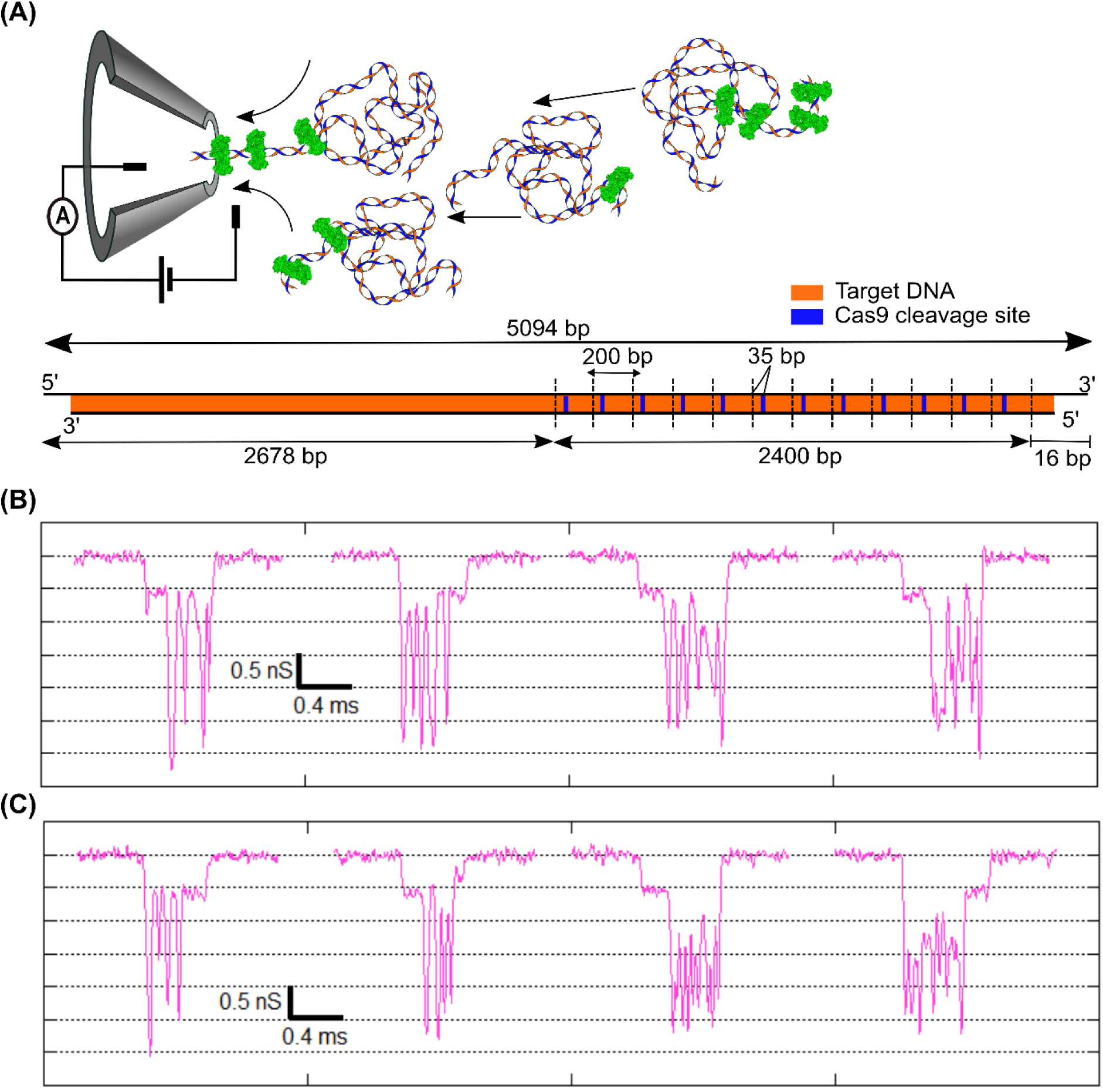
Detection of complex with multiple Cas9 bound to the DNA. (A) Schematic of the multi-Cas9 bound DNA sample and the translocation process through the nanopore. Multiple Cas9 protein bound on DNA is shown in green. (Bottom) shows the 5094 bp DNA (5 kb, in orange) with 12 repeats of 200 bp region (dotted vertical lines), each carrying a Cas9 target site shown in blue. In our experimental conditions, Cas9 to DNA ratio is chosen such that sub-saturated (3-5 Cas9 bound per DNA) arrays are formed. (B) and (C) show a library of representative events demonstrating multiple Cas9 protein bound on the DNA being detected as multiple spikes in events recorded during translocation of individual complex molecules.

To further investigate the readout of protein position on DNA from the spike position in the translocation event, we prepared a new Cas9-DNA complex with its target location shifted to the edge (215 bp from the start of the DNA) of the linear DNA (see schematic in Figure 3C). This sample is called edge-target in the text. Sample with edge target was measured on the same nanopore (as the center-target sample) and the events are shown in Figure 3C (right) and Figure S8D. Here we again see the bound protein as current spikes on top of the DNA blockade events. However, the location of the protein spike is now moved either to the beginning or the end of the event length. These two states again demonstrate that the protein carrying DNA may enter the nanopore in either orientation which brings the protein at either the start or the end of the event. Note, the protein spikes are also present for Cas9-bound DNA molecules that translocate in (partially or fully) folded conformation. Some of the representative events are shown in Figure S9 and S10 (see also Table S4).

We analyze the events with well-defined protein spikes to localize physical position of the protein on the DNA. We employ our, previously established, ECD based analysis method to correlate the position of electrical spikes found in translocation events to the actual location of Cas9 protein bound on the linearly translocating DNA^5^. Note that for this analysis only those events are considered where the DNA translocated in unfolded configuration. This was done to eliminate challenges brought in by events with spikes on folded or partially folded DNA. We calculated the relative location of Cas9 spike by taking ratio of the ECD up to the center of protein spike to the total ECD of the event, as shown below:

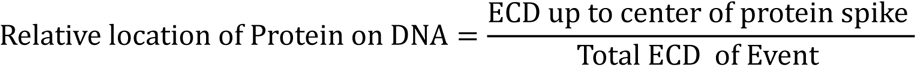

Spike location, relative to the DNA event (event length scaled to 1) was calculated using the above expression for each event. Figure 3B shows the histogram of identified spike locations relative to DNA length, for center-target sample. The two distinct peak positions (corresponding to the two entry orientations) were found to be 0.38 ± 0.05 and 0.61 ± 0.06. This result was reproduced on multiple nanopores and sample preparations, as seen in Table S5. From our experiments, the average value of the protein positions for center-target sample, were found to be 0.4 ± 0.01 and 0.59 ± 0.01, respectively. Our results show excellent agreement with true protein positions (0.39 and 0.61, see Figure 3A) of Cas9 bound to its targets with < 4 % error (see Table S5).

We reproduce the same analysis for the edge-target sample. Figure 3D shows the histogram of the protein spike positions measured in translocation data. We find that the mean values of peak location were found to be at 0.15 ± 0.06 and 0.86 ± 0.05, corresponding to the two orientations the protein-DNA complex enters the nanopore. This is also repeated on multiple nanopores as shown in Table S5 and Figure S11. The mean values of the relative protein location were estimated to be 0.17 ± 0.05 and 0.85 ± 0.03, respectively. The true values of the relative location for the edge-target sample is calculated to be 0.08 and 0.92. In the case of edge target sample, the protein location is at the very edge of the DNA. The observed deviation in measured protein location, in this case, is possibly due to large velocity fluctuation at the entry and exit of translocation as reported earlier^49–51^. From Figure 3B and 3D, we also note that the probability for entry of either end of the DNA into the nanopore was similar in both center and edge-target sample. This suggest that the Cas9 bound DNA molecule randomly selects, from the well equilibrated ensemble of both orientations, any one end to enter into the nanopore, as reported earlier^52^.

The ECD of the protein spikes in the translocation events also allows us to estimate the hydrodynamic volume of the Cas9 protein. First, we extracted the protein spike for every event with a custom written LabVIEW code and calculated its translocation characteristics, such as its ΔG, Δt and the ECD (see Figure S7 & Table S6). The power law dependence of ECD on the volume of the translocating molecule was measured earlier in Figure 1E (inset). By using volume of the 3 kb DNA, measured on the same nanopore, as *in-situ* control, we estimate the volume of Cas9 protein from multiple nanopores (see Table S6). From our experiments, the average volume of the Cas9 protein was found to be 3336.1 ± 388 nm^3^. Our measured protein volume agrees with the literature^28^ value of 3057.6 nm^3^, with less than 10 % error (see Table S6).

### Detection of DNA-Cas9 multi-protein complex

Next, we expand application of our method to demonstrate detection of multiple Cas9 proteins bound on the same DNA molecule. For this experiment we used 5094 bp (∼5 kb) linear DNA with 12 repeats of the 200 bp region with one Cas9 target per region. The Cas9 target is located 35 bp from the beginning of each 200 bp region. Schematic of the DNA with multiple target sites is shown in Figure 4A (right). In this experiment, we optimized the DNA and crRNA loaded Cas9 protein ratio so as to create sub-saturated (3 – 5 Cas9 protein per DNA) binding of Cas9 on the 12 target sequences. This ratio was optimized to ensure clear identification of multi-peaks during translocation signifying multiple proteins bound on the DNA. This was found to be essential for reproducibility of the results (as higher concentration of protein would clog the nanopore prematurely) as well. The multi-Cas9-DNA complex were translocated through the 20 nm nanopore at 300mV. A collection of representative events showing translocation signal for the sample is shown in Figure 4B and 4C. The events show DNA translocation with multiple protein spikes distributed along the DNA level. We attribute these spikes to multiple Cas9 bound to target DNA, since they have similar characteristics as shown with single Cas9-DNA complexes (see Figure 2F). Since the targets are located on one side of the DNA and the translocation could happen in either orientation, we find that the protein spikes were distributed randomly at either the start or end of event, as can be seen in Figure 4B and 4C. In multiple repeats of our experiments, we observe 20 -25 % of events to be over the ΔG threshold (see Figure 2E) that correspond to Cas9-DNA complexes, of which 50 -70 % of events show two or more spikes per event. We observed a variety of events with anywhere between 1 to 6 protein spikes. A library of such events is shown in Figures S12. In addition to the unfolded events, Cas9-DNA complex molecules can translocate in folded conformation producing complicated events structures. Some of such representative events are shown in Figure S13A & S13B. These results and event characteristics were reproduced on different nanopores as tabulated in Table S7.

### Characterization of enzymatic activity of catalytically active Cas9

We next apply the robust nature of the ECD-based analysis, to quantify enzymatic activity of the wild-type DNA-binding enzyme, Cas9. It is shown earlier that the wild-type Cas9 enzyme remains stably bound to its target DNA until the cleaved products are released upon exposure to agents, such as salt, temperature or detergent^33^. We quantify time-dependent release of the cleaved products to measure the temporal stability of catalytically active Cas9 enzyme on the target DNA, in our experimental conditions.

To measure the constituents of our Cas9-DNA complex sample, we calculated the ECD of every collected event. Schematic of this experiment is shown in Figure 5A. The histogram of the estimated ECD values is shown in Figure 5C (top). We find four distinct peaks in the ECD histogram, which should correspond to four distinct populations. Gaussian fitting of the four peaks reveal the mean values as ECD_1_ = 59.46 ± 9.46 (ke), ECD_2_ = 106 ± 10.65 (ke), ECD_3_ = 184.16 ± 12.95 (ke) and ECD_4_ = 239.39 ± 49.44 (ke). These values are also shown as vertical dotted lines for visual clarity.

**Figure 5:**
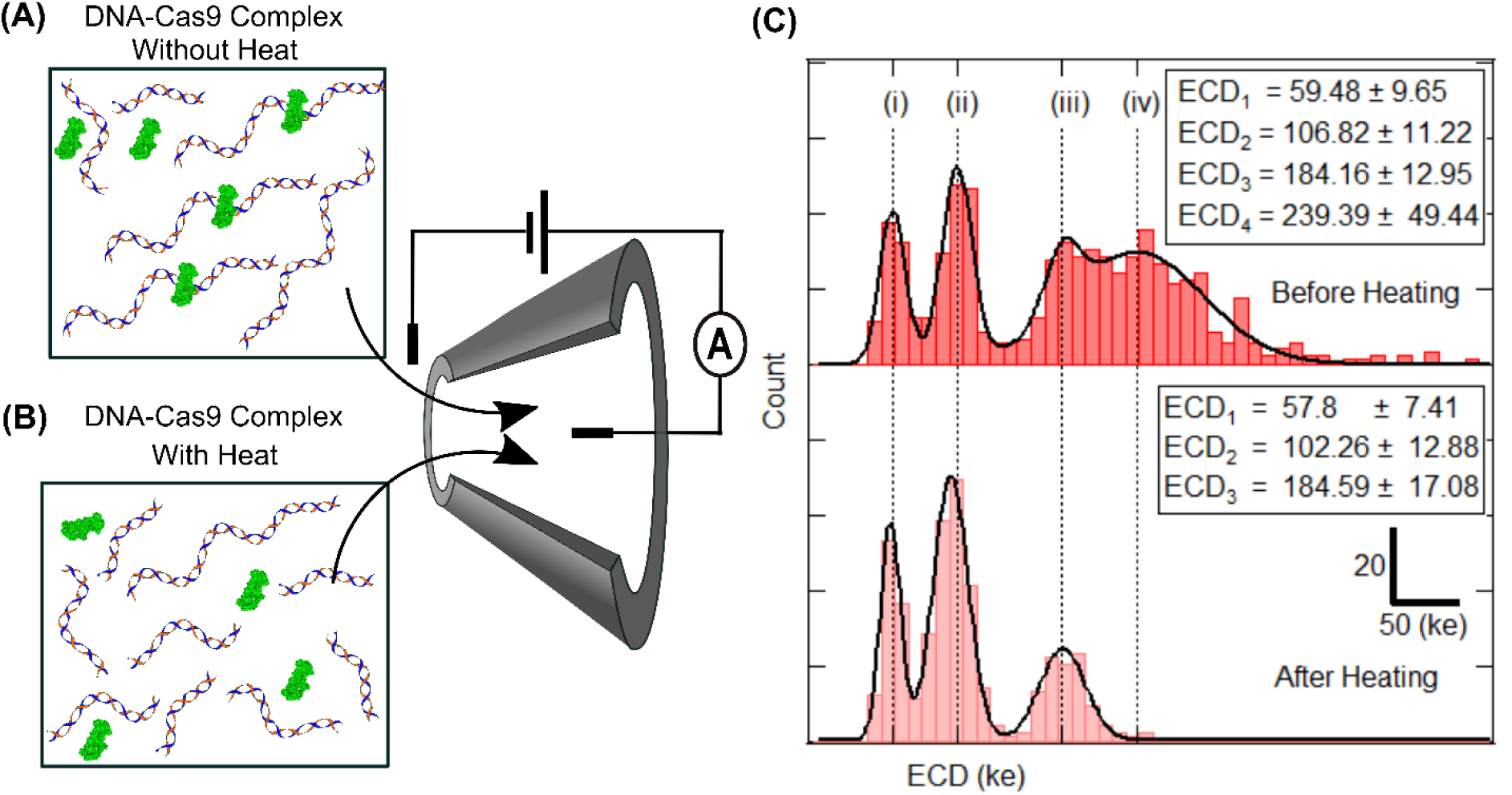
Detection of Cas9-bound and Cas9-cleaved DNA. (A) shows cartoon of Cas9-DNA complex sample at room temperature. The reaction mix contains free DNA, free Cas9 protein, Cas9-bound DNA and Cas9-cleaved DNA fragments. (B) shows cartoon of the Cas9-DNA complex sample after heat-release of products. Sample contains denatured Cas9 protein, released cleaved DNA products and free DNA. Both samples are measured using the same nanopore, back-to-back, as shown in the middle schematic. (C) shows ECD histograms of the two, room temperature (top) and heat-released (bottom), samples. Histograms show populations for Cas9-bound DNA (iv), free DNA (iii) and the two Cas9-cleaved products (ii) and (i), as distinct ECD peaks. Note the absence of Cas9-bound DNA peak in the heat-released sample.

**Figure 6:**
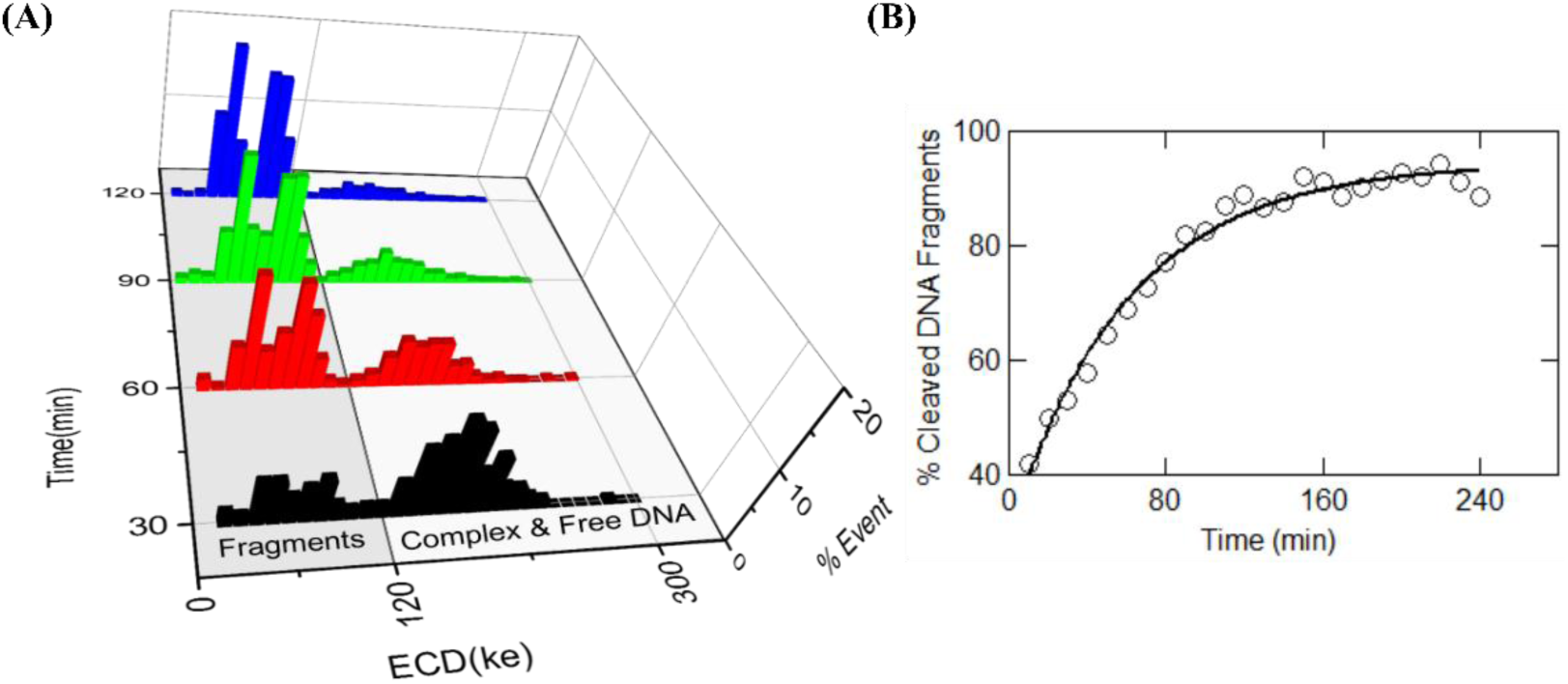
Real-time release of cleaved products by catalytically active wild-type Cas9 protein. (A) quantifies different populations of DNA fragments that are released by Cas9 cleavage at room temp. Events are accumulated for 30-min time intervals and shown at time points of 30, 60, 120 and 180 min (measured at 500 mV). Note the decrease in Cas9-bound DNA population and increase in the cleaved DNA fragments population with time. The percentage of cleaved fragments was calculated from the number of events under the ECD histograms (shaded region in (A)) collected for each 10 min interval and is plotted in (B). The solid line shows fit to a first order rate equation with rate constant of 0.017 ± 0.0013 min^−1^.

To confirm the identities of our ECD peaks, we re-measured translocation of the same sample, after heat-release of the products, in the same nanopore. The heat treatment of the Cas9-DNA complex sample at 90° C for 10 min denatures the Cas9 protein resulting in complete release of the cleaved fragments (see Figure 5B and Figure S1 (lane 13 in (A)). In the ECD histogram of the heat-treated sample, we observe only three distinct peaks (see Figure 5C (Bottom)). Comparing the locations of the peaks in the two histograms (Figure 5C (top & bottom)), we find that the population with the largest ECD value (population (iv)) is absent, whereas the other three peaks remain as is. This confirms our annotation of the population marked (iv) to be the Cas9-bound DNA complex and population (iii) as the Cas9-free target DNA. Consequently, the peaks (i) and (ii) are identified to be the populations corresponding to the two cleaved fragments that are released after Cas9 cleavage of the target DNA. Note, ECD histograms of the translocation of control DNA sample do not show multiple peaks (see Figure S15). These results were reproduced in multiple nanopores and multiple sample preparations (see Figure S14 and Table S8).

We quantify the length of the released DNA fragments (population (i) and (ii)), from the ECD peak values using the power law dependence established earlier in Figure 1E inset. Using the ECD value of the free DNA (3 kb) (shown in Figure 5C (bottom)) as the *in situ* control, we estimate the DNA lengths of the populations (i) and (ii) using the expression 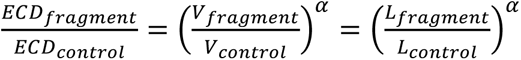. From the above calculations the lengths of the DNA fragments in population (i) and (ii) were estimated to be 1122 bp and 1826 bp respectively. The expected value of the two fragments are 1166 bp and 1859 bp. Our estimated DNA fragment length values, measured in multiple nanopores, agree with the expected fragment lengths with-in 4 % error (see Table S8A).

This allows us to demonstrate, for the first time, simultaneous detection and quantification of enzyme-bound, enzyme-free and enzymatically cleaved product populations, in the nanopore platform.

Above quantification of the components of a DNA-protein reaction mixture also allows us to measure the temporal progression of Cas9 based cleavage of the target DNA. For this experiment we continuously recorded the translocation events of the Cas9-DNA complex sample, through the nanopore (at 500 mV), for up to 4 hours. At different time points, we quantified relative fractions of the cleaved products using the population identification strategy developed by ECD analysis. Figure 6A shows ECD histograms of events collected at four different time points. It shows peaks corresponding to the two cleaved fragments (shaded region) and the population corresponding to the free and Cas9-bound DNA (unshaded region). Note that due to very small ECD differences between the third and the fourth peaks, they sometimes are indistinguishable. The two peaks corresponding to the cleaved fragments are, however, always resolvable and the number of events in these two peaks increase with time, with corresponding decrease in the higher ECD peaks. This indicates time dependent release of cleaved products from the Cas9-bound DNA complexes. We calculated the percentage of cleaved products, by counting the number of events under the cleaved products ECD peaks, at every 10 min interval and show this progression in Figure 6B. The percentage of cleaved fragments increased exponentially with time and saturate after about 180 min. We fitted the first order rate equation to the data in Figure 6B and find the rate of release of the cleaved products to be 0.017 ± 0.0013 min^−1^. We produced these results in multiple nanopores. We also found that the product release rates are intrinsic to the Cas9 enzyme and are independent of the voltage applied across the nanopore (see Figure S16 and Table S9).

## CONCLUSION

In this work we have presented a nanopore-based assay to localize Cas9 bound at different locations on the target DNA. We show that the Cas9-bound DNA molecules can be uniquely identified from the unbound background by a high-resolution secondary current spike during the translocation event of the complexed molecule. A detailed ECD-based analysis was employed to identify the location of the centrally-bound Cas9 protein on the DNA with 4 % accuracy. This analysis also allowed us to quantify the volume of the bound protein within 10% error of the literature value. For the Cas9-DNA complex, excellent correlation of the location of the current spikes in the translocation events to the physical location of the protein on the DNA was established for Cas9 positioned at multiple places. We employed this detection scheme on DNA complexed with multiple (1 – 6) Cas9 proteins which showed corresponding multi-peaks in the translocation events and allowed us to count the number of bound proteins. Finally, we showed the quantification of cleaved and released products of the catalytically active Cas9. The unique sizes of the product fragments were quantified using their unique ECD signatures and the fragment lengths were quantified within 4 % error. We also show quantification of the rate of product release, with time, by the enzyme (in our experimental conditions). we find the release rate of Cas9 cleaved products to be 0.017 ± 0.0013 min^−1^. The experimental result presented has potential application for glass nanopore in Cas9 based targeting in genome engineering applications.

In conclusion, we have developed, through a range of experiments, a nanopore assay to fingerprint location as well as monitor functional activity of enzymes, such as endonucleases, restriction enzymes or any other DNA-binding operator protein, with single molecule resolution. Our nanopore sensor distinguishes template DNA when translocating bare as compared to when an enzyme molecule is bound to it. On a 3 kb template DNA we targeted binding of the Cas9 enzyme using crRNA. Nanopore events for bare DNA, with current level corresponding to DNA translocation, could be isolated from the events for the enzyme bound DNA solely based on the high-amplitude secondary spike in current on top of the DNA level. These secondary spikes were shown to be due to the enzyme bound on the DNA. We showed that the relative location of the spike in the event correspond to relative location of the enzyme on the DNA. This fingerprinting of protein location was further confirmed by targeting the Cas9 protein to a 2^nd^ location and measuring corresponding shift in the relative spike location. We show that ECD-based analysis is a powerful technique to quantify protein position as well as protein volume from these protein spikes. Importantly, the assay presented here allowed us to directly measure, in real-time, the catalytic output of DNA fragments resulting from active Cas9 cleavage. This combination of high signal-to-noise combined with ECD analysis of events resulting in real-time analytical quantitation of enzyme reaction output opens up the field to study DNA and/or RNA specific binding proteins, protein complexes bound to DNA as well as chromatin structure.

## ACKNOWLEDGEMENTS

We acknowledge RRI’s internal funding and assistance from Mr. Yatheendran & RRI SEM facility in this work. We also acknowledge SJRI administration for permissions.

## AUTHORSHIP CONTRIBUTIONS

GVS, and PN conceived the project. PN & GVS planned the experimental protocol. DS & PN contributed in sample preparation. PN and SS performed experiments. PN, SS and GVS performed data analysis. All authors contributed in manuscript writing.

