## Supplementary File for "Nanopore Assay for Fingerprinting DNA Binding and Quantifying Real-Time Cleavage by Catalytically Active Cas9 Enzyme"

Punitkumar Nagpure, Sarangi Suresh, Divya Shet, and Gautam V. Soni
Raman Research Institute, Bangalore – 560080 INDIA

### Supplementary Information file

This PDF file includes:
Tables S1-S9
Figures S1-S16

| DNA size in kbp | Concentration (nM) Pore 1 | Concentration (nM) Pore 2 | Concentration (nM) Pore 3 |
|---|---|---|---|
| 0.5 | 0.53 | 1.1 | 0.53 |
| 1 | 0.27 | 0.53 | 0.27 |
| 1.5 | 0.15 | 0.3 | 0.15 |
| 2 | 0.15 | 0.3 | 0.15 |
| 3 | 0.26 | 0.48 | 0.26 |
| 4 | 0.05 | 0.1 | 0.05 |
| 5 | **0.43** | **0.85** | **0.43** |
| 6 | 0.05 | 0.11 | 0.05 |
| 8 | 0.03 | 0.07 | 0.03 |
| 10 | **0.38** | **0.77** | **0.38** |

***Table S1: Experimental concentration of custom mix –DNA sample:*** *The table shows concentration of each DNA length (from manufacturer's data sheet) in our custom mix DNA sample used in three different experiments. The 5 and 10 kb DNA (shown in bold) were added externally to the 1 kb DNA ladder.*

| DNA (kbp) | ECD peak Pore 1 | ECD peaks Pore 2 | ECD peaks Pore 3 |
|---|---|---|---|
| 0.5 | 24.6 ± 5.35 | 28.71 ± 7 | 24.48 ± 7.21 |
| 1 | 52 ± 7.63 | 64.13 ± 9.3 | 52.8 ± 9.71 |
| 1.5 | 83.1 ± 11.05 | 101.65 ± 12.04 | 86.28 ± 13.08 |
| 2 | 117.8 ± 13.17 | 144.54 ± 16.6 | 122.67 ± 14.37 |
| 3 | 196 ± 28.02 | 238.48 ± 32.39 | 197.17 ± 23.37 |
| 5 | 358.5 ± 39.23 | 434.56 ± 52.28 | 361.91 ± 68.61 |
| 10 | 762.9 ± 97.07 | 915.05 ± 106.59 | 789.39 ± 93.69 |

***Table S2: Summary of ECD peak values measured for custom mix DNA sample:*** *The table shows ECD peak fit values for each length of DNA, measured in three independent experiments (see Figure 1E & Figure S3). Std deviation of ECD histograms and the power-law relationship between ECD (ke) and DNA length (kb), was used to estimate resolution of DNA length measured to be 189 bp for DNA lengths of < 2kbp.*

| Pore ID | A | α |
|---|---|---|
| 1 | 0.003 ± 0.0017 | 1.17 ± 0.06 |
| 2 | 0.0039 ± 0.0021 | 1.16 ± 0.06 |
| 3 | 0.0031 ± 0.002 | 1.17 ± 0.07 |
| Mean | 0.0033 ± 0.0011 | 1.17 ± 0.04 |

***Table S3: Summary of power-law fitting to ECD vs volume for translocation custom mix DNA sample:*** *The table shows fitting parameter values of A and α obtained by fitting (ECD (ke) = A × $(V_{DNA})^{\alpha}$), to the ECD vs DNA volume data. Reproducibility from three different nanopore experiments is shown.*

| Pore ID | Sample | Total Events recorded | % Cas9-DNA Complex | % Cleaved Products | % unfolded complex Event | % folded Complex Events |
|---|---|---|---|---|---|---|
| 4 | Control DNA | 1567 | | | 44 | 56 |
| | Cas9-DNA Complex (Center Target) | 589 | 44.14 | 15 | 54 | 46 |
| | Cas9-DNA Complex (Edge Target) | 915 | 28.3 | | 35 | 65 |
| 5 | Control DNA | 1132 | | | 33.39 | 66.6 |
| | Cas9-DNA Complex (Center Target) | 929 | 48.76 | 26 | 40.17 | 59.82 |
| | Cas9-DNA Complex (Edge Target) | 1133 | 36.98 | | 30 | 70 |
| 6 | Control DNA | 864 | | | 41 | 59 |
| | Cas9-DNA Complex (Center Target) | 672 | 66.66 | 20 | 31 | 69 |
| | Cas9-DNA Complex (Edge Target) | 845 | 58.69 | | 25 | 75 |
| 7 | Control DNA | 1421 | | | 43 | 60 |
| | Cas9-DNA Complex (Center Target) | 764 | 60 | 32 | 40.6 | 59.4 |
| 8 | Control DNA | 1178 | | | 37 | 63 |
| | Cas9-DNA Complex (Center Target) | 753 | 53.65 | 31 | 40 | 60 |
| 9 | Control DNA | 1112 | | | 55.67 | 44.33 |
| | Cas9-DNA Complex (Center Target) | 922 | 33.51 | 49.89 | 54.37 | 45.63 |
| 10 | Control DNA | 1130 | | | 37.08 | 62.92 |
| | Cas9-DNA Complex (Center Target) | 537 | 51.58 | 39.11 | 46.21 | 53.79 |
| 11 | Control DNA | 1260 | | | 44.84 | 55.16 |
| | Cas9-DNA Complex (Center Target) | 1027 | 41.09 | 55.4 | 44.79 | 55.21 |
| 12 | Control DNA | 1417 | | | 51.24 | 48.76 |
| | Cas9-DNA Complex (Center Target) | 536 | 50.56 | 39.74 | 59.04 | 40.96 |
| 13 | Control DNA | 959 | | | 43.59 | 56.41 |
| | Cas9-DNA Complex (Center Target) | 408 | 64.46 | 25 | 56.27 | 43.73 |
| 14 | Control DNA | 1380 | | | 50.14 | 49.86 |
| | Cas9-DNA Complex (Center Target) | 3817 | 30.97 | 48.05 | 54.48 | 45.52 |
| 15 | Control DNA | 1826 | | | 50.49 | 49.51 |
| | Cas9-DNA Complex (Center Target) | 914 | 50.11 | 45.4 | 46.72 | 53.28 |
| 16 | Control DNA | 1511 | | | 49.77 | 50.23 |
| | Cas9-DNA Complex (Center Target) | 2031 | 29.54 | 72.77 | 49.83 | 50.17 |
| 17 | Control DNA | 1193 | | | 52.39 | 47.61 |
| | Cas9-DNA Complex (Center Target) | 752 | 45.61 | 47.21 | 66.18 | 33.82 |
| 18 | Control DNA | 1112 | | | 46.58 | 53.42 |
| | Cas9-DNA Complex (Center Target) | 918 | 47.06 | 30.83 | 77.55 | 22.45 |

***Table S4: Summary of Cas9-DNA complex events.*** *Percentage of bound, cleaved, folded and linear conformation, as measured from ECD analysis, is shown. For pore IDs 4,5 and 6, all three samples were compared on the same nanopore, back-to-back. The results were reproduced on multiple nanopores, as shown.*

| Sr/No | Pore ID | Sample | Total Number of events Selected | Scaled position of spike on DNA event | |
|---|---|---|---|---|---|
| | | | | $Z_1$ | $Z_2$ |
| 1 | 4 | Cas9-DNA Complex (Center Target) | 141 | 0.41 ± 0.03 | 0.58 ± 0.06 |
| | | Cas9-DNA Complex (Edge Target) | 91 | 0.17 ± 0.11 | 0.82 ± 0.04 |
| 2 | 5 | Cas9-DNA Complex (Center Target) | 182 | 0.41 ± 0.05 | 0.6 ± 0.05 |
| | | Cas9-DNA Complex (Edge Target) | 111 | 0.18 ± 0.09 | 0.86 ± 0.04 |
| 3 | 6 | Cas9-DNA Complex (Center Target) | 138 | 0.38 ± 0.05 | 0.61 ± 0.06 |
| | | Cas9-DNA Complex (Edge Target) | 121 | 0.15 ± 0.06 | 0.86 ± 0.05 |
| 4 | 7 | Cas9-DNA Complex (Center Target) | 186 | 0.39 ± 0.06 | 0.59 ± 0.05 |
| 5 | 8 | Cas9-DNA Complex (Center Target) | 160 | 0.39 ± 0.05 | 0.6 ± 0.05 |
| 6 | 9 | Cas9-DNA Complex (Center Target) | 168 | 0.40 ± 0.04 | 0.58 ± 0.04 |
| 7 | 10 | Cas9-DNA Complex (Center Target) | 128 | 0.39 ± 0.04 | 0.59 ± 0.05 |
| 8 | 11 | Cas9-DNA Complex (Center Target) | 189 | 0.4 ± 0.05 | 0.6 ± 0.05 |
| 9 | 12 | Cas9-DNA Complex (Center Target) | 160 | 0.39 ± 0.04 | 0.6 ± 0.05 |
| 10 | 13 | Cas9-DNA Complex (Center Target) | 148 | 0.41 ± 0.05 | 0.59 ± 0.05 |
| 11 | 14 | Cas9-DNA Complex (Center Target) | 644 | 0.40 ± 0.05 | 0.60 ± 0.05 |
| 12 | 15 | Cas9-DNA Complex (Center Target) | 214 | 0.42 ± 0.05 | 0.60 ± 0.03 |
| 13 | 16 | Cas9-DNA Complex (Center Target) | 299 | 0.39 ± 0.04 | 0.58 ± 0.05 |
| 14 | 17 | Cas9-DNA Complex (Center Target) | 227 | 0.39 ± 0.06 | 0.59 ± 0.04 |
| 15 | 18 | Cas9-DNA Complex (Center Target) | 335 | 0.39 ± 0.06 | 0.59 ± 0.05 |
| Mean | | Cas9-DNA Complex (Center Target) | | 0.40 ± 0.01 | 0.59 ± 0.01 |
| error | | Actual relative positions Z1 = 0.39 and Z2 = 0.61 | | 2.56 % | 3.39 % |
| Mean | | Cas9-DNA Complex (Edge Target) | | 0.17 ± 0.05 | 0.85 ± 0.03 |
| error | | Actual relative positions Z1 = 0.08 and Z2 = 0.92 | | 112 % | 7.6 % |

***Table S5: Summary of location estimation of Cas9 bound on 3 kb DNA:*** *The table shows the estimated location of Cas9 protein bound on center and/or edge target DNA measured with multiple nanopores. The experiment was repeated in 15 different pores of similar size. Mean positions and percentage errors in position localization (for both center & edge target) are shown in last 4 rows.*

| PoreID | Control DNA (3 kb) ECD ± SD (ke) | Spike ECD (Cas9) Mean ± SD (ke) | Volume of Cas9 Protein ($nm^3$) $\frac{ECD_{Cas9}}{ECD_{3\ kb}} = \left(\frac{V_{Cas9}}{V_{3\ kb}}\right)^{\alpha}$ |
|---|---|---|---|
| 4 | 215.02 ± 22.81 | 39.29 ± 18.38 | 3021.74 |
| 5 | 218 ± 23.9 | 43.03 ± 17.57 | 3227.74 |
| 6 | 231.69 ± 22.74 | 48.91 ± 28.26 | 3418.50 |
| 7 | 172.36 ± 17.21 | 48.32 ± 27.49 | 4356.47 |
| 10 | 196.15 ± 20.71 | 40.77 ± 20.73 | 3373.47 |
| 11 | 227.34 ± 22.18 | 46.42 ± 26.80 | 3322.58 |
| 8 | 255.08 ± 27.33 | 45.94 ± 20.21 | 3040.01 |
| 12 | 153.28 ± 16.15 | 30.16 ± 16.11 | 3219.06 |
| 15 | 153.16 ± 15.18 | 33.35 ± 28.49 | 3510.26 |
| 13 | 233.28 ± 24.19 | 40.22 ± 23.48 | 2875.31 |
| | Mean ± SD | | 3336.51 ± 387.83 $nm^3$ |
| | Literature Reported value | | 3057.6 $nm^3$ |
| | Relative Percentage Error | | 9.12 % |

***Table S6: Summary of ECD values of the control DNA and protein spikes:*** *The Table shows the summary of ECD value of control DNA and protein spike for complex translocation measured on the same nanopore. Volume of Cas9 protein is estimated from the ECD values of the spikes by comparing it to the known volume of the control (3 kb) DNA. Last row shows the relative error in estimated volume of Cas9 protein to be less than 10 % of the literature value (see reference 28 in main text).*

| Pore ID | Total Events | #events above threshold | % of events with # of spikes < 2 | % of events with # of spikes >=2 | % of unfolded events | % of folded events |
|---|---|---|---|---|---|---|
| 22 | 2073 | 306 | 15 | 37 | 78 | 22 |
| 23 | 659 | 161 | 24 | 29 | 55 | 45 |
| 24 | 2069 | 560 | 27 | 36 | 68 | 32 |
| 25 | 1067 | 268 | 25 | 47 | 55 | 45 |
| 26 | 2196 | 495 | 22 | 77 | 57 | 43 |
| 27 | 2490 | 619 | 24 | 57 | 66 | 34 |
| 28 | 903 | 267 | 29 | 53 | 74 | 26 |
| 29 | 934 | 255 | 27 | 55 | 54 | 46 |
| 30 | 906 | 236 | 26 | 74 | 75 | 25 |

***Table S7: Summary of events showing multi-Cas9 Translocation:*** *The table shows statistics of multi-spike events for translocation of complex with multiple Cas9 proteins bound to the DNA. The results were reproduced in multiple experiment performed on similar sized nanopores.*

| Pore ID | Total Events | Fragment - 1 | | Fragment - 2 | | Free DNA | | Length Fragment-1 | Length Fragment-2 |
|---|---|---|---|---|---|---|---|---|---|
| | | ECD ± SD | % of events | ECD ± SD | % of events | ECD ± SD | % of events | | |
| 17 | 462 | 57.81 ± 7.41 | 26.41 | 102.26 ± 12.88 | 49.35 | 184.56 ± 17.08 | 24.24 | 1122 | 1826 |
| 18 | 752 | 44.47 ± 7.09 | 37.63 | 78.78 ± 9.58 | 55.98 | 142.28 ± 18.02 | 6.38 | 1119 | 1825 |
| 19 | 1313 | 46.6 ± 6.95 | 32.44 | 81.39 ± 11.6 | 28.33 | 146.46 ± 16.10 | 39.22 | 1137 | 1831 |

***Table S8A: Summary of cleaved DNA lengths:*** *The table shows estimated length for the cleaved DNA fragments released from Cas9-DNA complex measured in three independent experiments.*

| Pore ID | Measured Value | | Actual Value | | Relative Error | |
|---|---|---|---|---|---|---|
| | Fragment -1 | Fragment - 2 | Fragment - 1 | Fragment - 2 | Fragment - 1 | Fragment - 2 |
| 17 | 1122 | 1826 | 1166 | 1859 | 3.77 | 1.78 |
| 18 | 1119 | 1825 | 1166 | 1859 | 4.03 | 1.83 |
| 19 | 1137 | 1831 | 1166 | 1859 | 2.49 | 1.51 |

***Table S8B: Comparison of true & experimentally calculated DNA fragment length:*** *Table shows the relative error in estimation of the cleaved DNA fragment lengths, measured in three independent experiments.*

| Pore ID | Duration | Voltage (mV) | k | $D_{free}$ | $C_o$ |
|---|---|---|---|---|---|
| 31 | 4hr | 500 | 0.017 ± 0.0013 | 5.64 ± 1.13 | 64.29 ± 2.12 |
| 32 | 4hr | 500 | 0.020 ± 0.0025 | 9.03 ± 2.13 | 85.75 ± 5.69 |
| 33 | 4hr | 300 | 0.010 ± 0.0028 | 10.95 ± 6.09 | 67.56 ± 5.05 |
| 34 | 2hr | 300 | 0.016 ± 0.012 | 36.32 ± 12.30 | 40.49 ± 9.04 |
| 35 | 2hr | 900 | 0.017 ± 0.0038 | 40.55 ± 4.16 | 55.034 ± 3.46 |

***Table S9: Summary of rate kinetics for release of Cas9-DNA complex:*** *The table shows the rate constant (k) for release of DNA fragments from Cas9-DNAcomplex in our experimental conditions. Here $D_{free}$ and $C_o$ are fractions of Cas9-free and Cas9-bound population (see main text).*

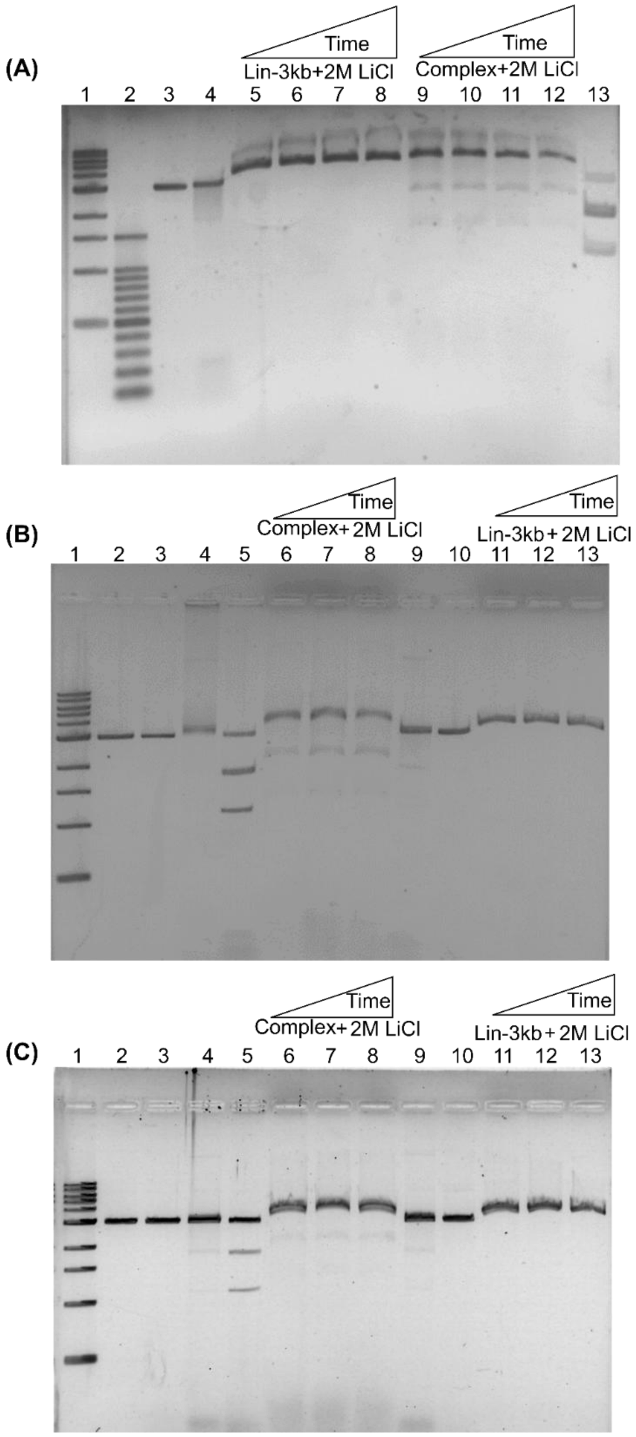


***Figure S1: Gel shift assay showing Cas9-DNA complex stability in experimental buffer.*** *(A) Lanes 1 & 2 are 1 kb and 100 bp ladder. Lanes 3 & 4 are free and Cas9-bound DNA sample in HEPES buffer. Control DNA (3025bp) (Lanes 5 – 8) and Cas9-DNA complex (Lanes 9 – 12) in salt for 30,60,90 & 120 min at room temperature, respectively. Note, presence of high salt in sample reduces band sharpness as well as shifts the bands slightly up. Lane-13 shows release of cleaved products upon heating the complex at 90 $^oC$ for 10 min. (B) and (C) shows reproducibility of result shown in (A). Lane -1: 1kb DNA ladder, Lane -2: Lin - 3kb (Control DNA), Lane -3: Lin 3kb DNA +Cas9 –RNA (Negative Control), Lane –4 & 9: Lin -3kb + Cas9 +RNA (Complex), Lane -5: Complex after heating at 90 $^oC$ for 10 min, Lanes 6 to 8: Cas9-DNA complex & Lanes 11 to 13: Control DNA in salt (2M LiCl) for 30, 60 and 90 min respectively.*

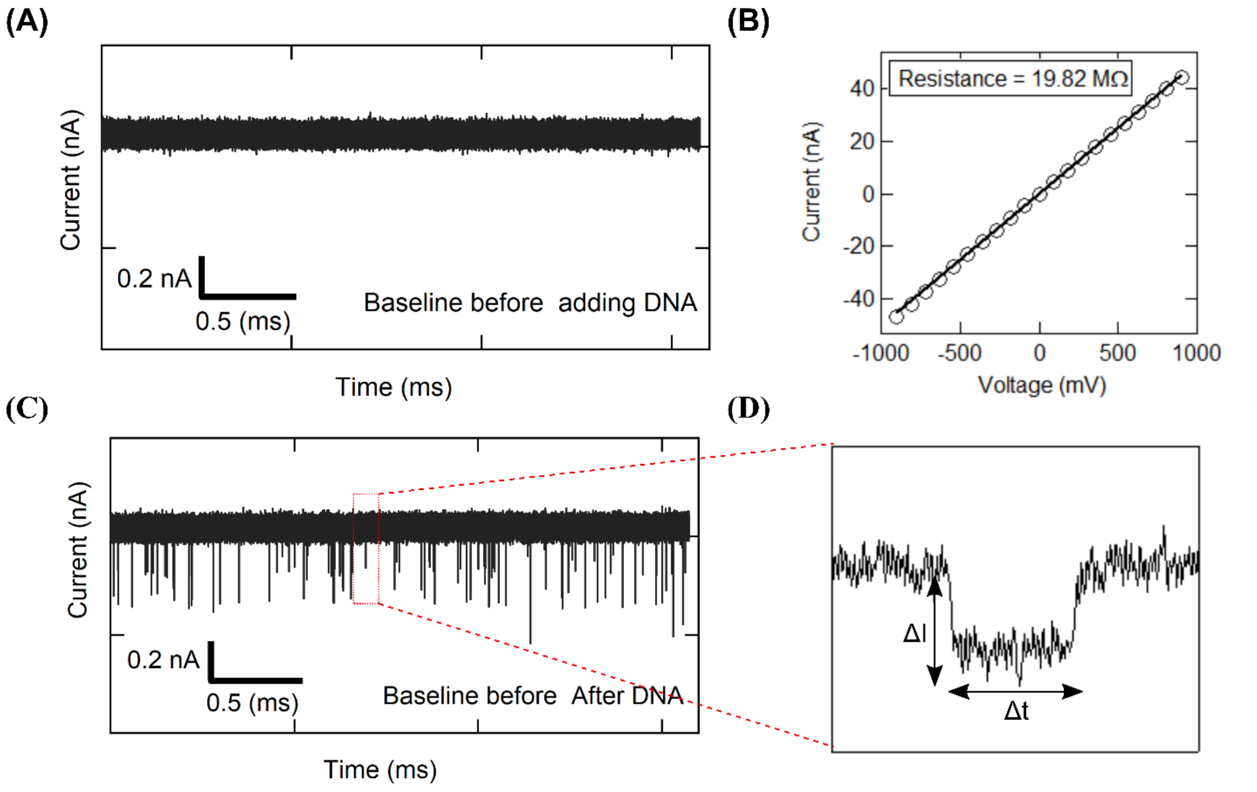


***Figure S2: baseline current signal of translocation of mixture of DNA.*** *(A) shows the baseline current through nanopore before addition of sample at 300mV. Before addition of sample baseline were stable and there were no spurious events. The pore was characterized by recording I-V curve as shown in (B). (C) Shows the events detected on the baseline current after addition of DNA sample. The zoom of one of these DNA translocation events is shown in (D), with current drop (ΔI) and dwell time (Δt) marked.*

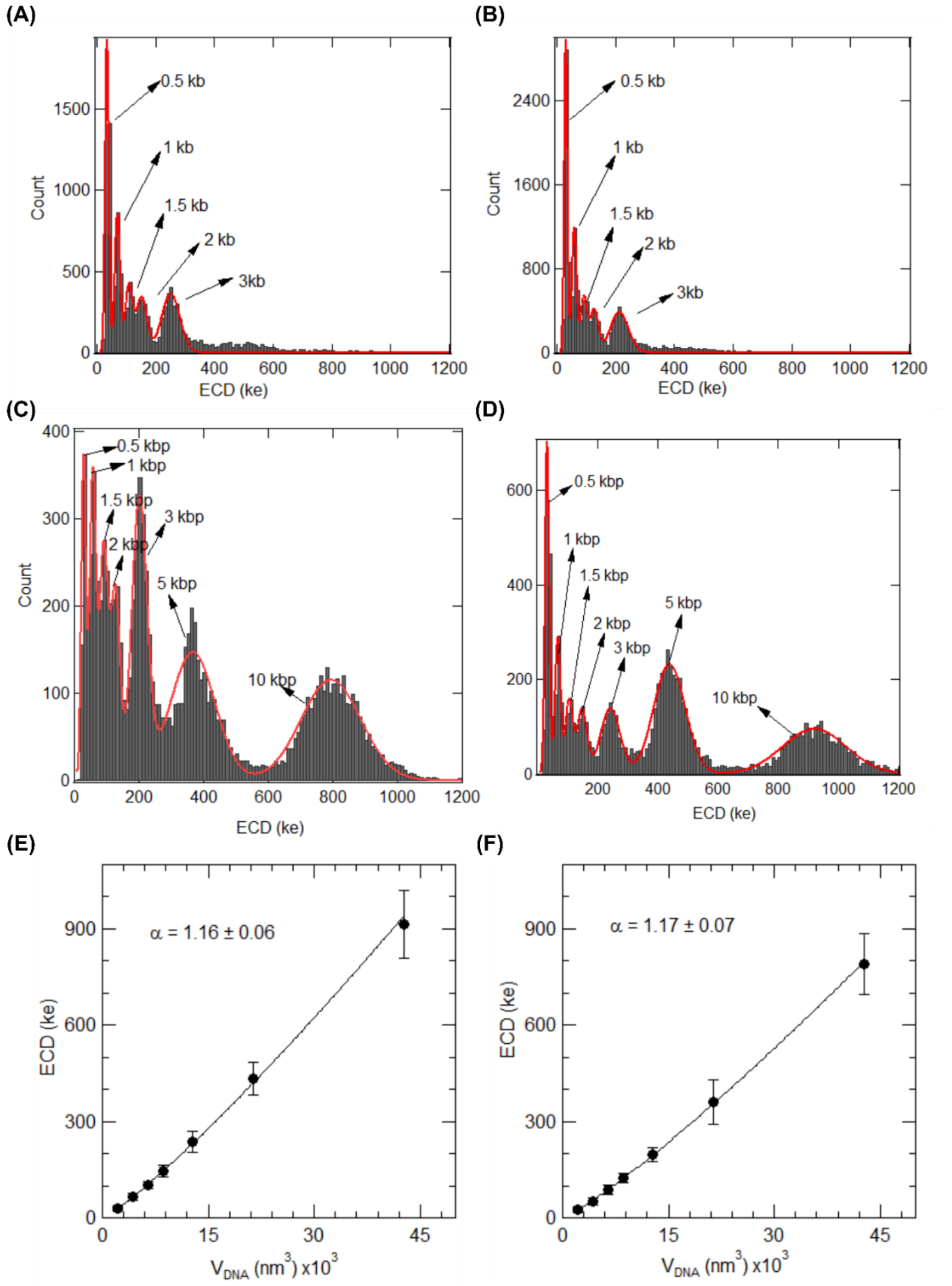


***Figure S3: ECD analysis of DNA molecules.*** *(A) and (B) shows ECD histograms of the events detected for of 1kb DNA ladder sample. Red solid lines are Gaussian fits to the peaks. (C) & (D) shows the ECD histograms (and Gaussian peak fits) of translocation events recorded for custom-mix of DNA lengths measured in two different nanopores of similar diameters. Custom-mix was made by adding 5 and 10 kb DNA to the 1 kb DNA ladder sample. (E) & (F) shows the relationship between the measured ECD peaks and the volume of the translocating DNA molecules in the custom-mix sample. The errorbars are calculated as width of Gaussian fits. Solid lines show the power-law fit (see text). The fit exponent is shown as well.*

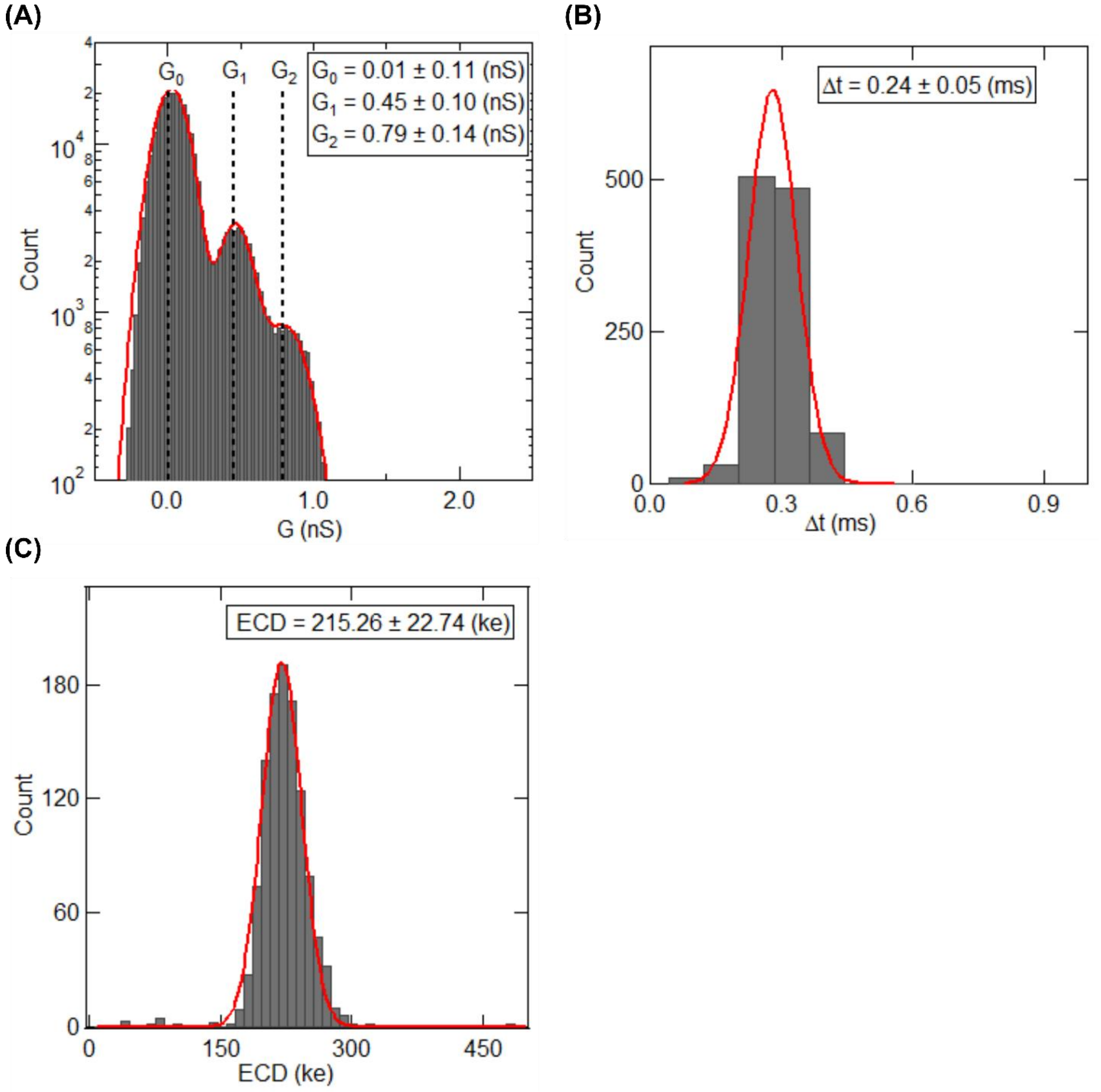


***Figure S4: Translocation characteristics of control DNA (3025 bp).*** *(A-C) shows the conductance, dwell time and ECD histograms of a typical 3 kb DNA translocation in a 20 nm nanopore.*

Calculation of $\Delta G$ values from conductance histogram: $G_0$, $G_1$ and $G_2$ represent mean values of the baseline, 1$^{st}$ (linear) and and 2$^{nd}$ (folded) blockade levels during DNA translocation.

$\Delta G_1 = G_1 – G_0 = 0.44 \pm 0.15$ (nS) (linear DNA)
$\Delta G_2 = G_2 – G_0 = 0.78 \pm 0.18$ (nS) (Folded DNA)

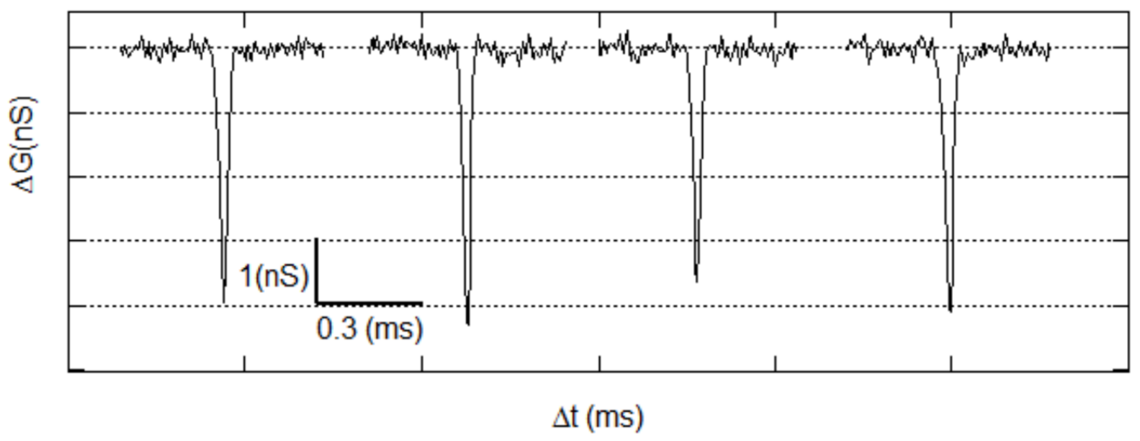


***Figure S5: Representative events for control DNA (3025 bp).*** *shows examples of 3 kb translocation events (less than 0.05 % of total events) where the event depth crosses the chosen threshold ($\Delta G > 1.2$ nS) for spike detection.*

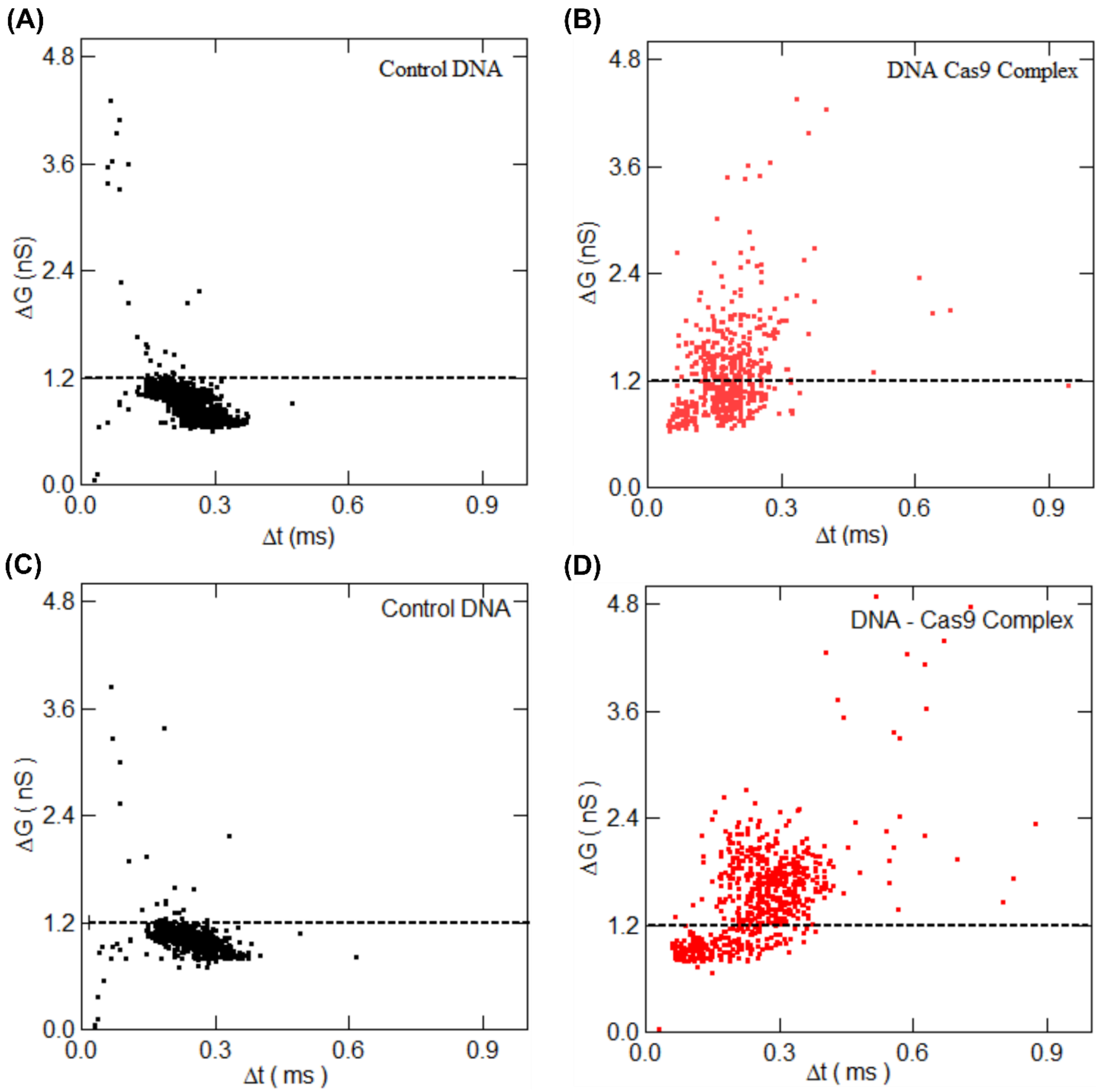


***Figure S6: Detection of Cas9-DNA complex measured in different nanopores (300mV).*** *Shows reproducibility of data shown in Figure 2. (A & C) show the scatter plot of conductance vs dwell time of control DNA and (B & D) shows the same scatter plot for Cas9-DNA complex recorded in poreIDs 5 & 6, respectively.*

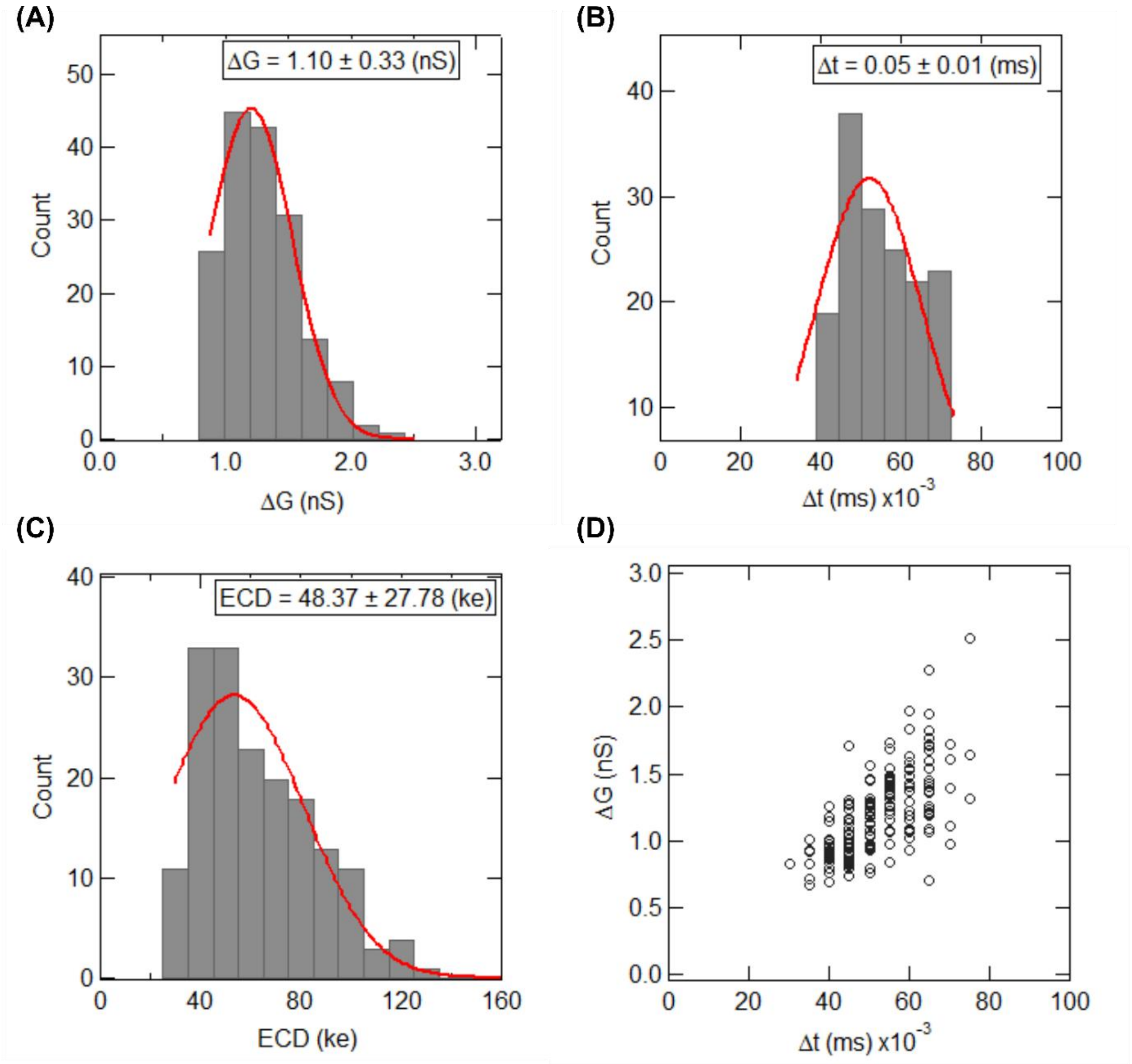


***Figure S7: Translocation characteristics of bound Cas9 on DNA.*** *(A), (B) & (C) shows the typical histogram of ΔG, Δt and ECD of Cas9 protein spike extracted from Cas9-DNAcomplex events. Figure (D) represent scatter plot of ΔG Vs Δt.*

Signal of Protein bound DNA:

$\Delta G_{DNA+protein} = \Delta G_{DNA} + \Delta G_{protein\ spike} = 0.46 + 1.1 = 1.56$ (nS)

$$Signal\ of\ protein\ bound\ on\ DNA = \frac{\Delta G_{DNA+protein}}{\Delta G_{DNA}} = \frac{1.56}{0.46} = 3.4$$

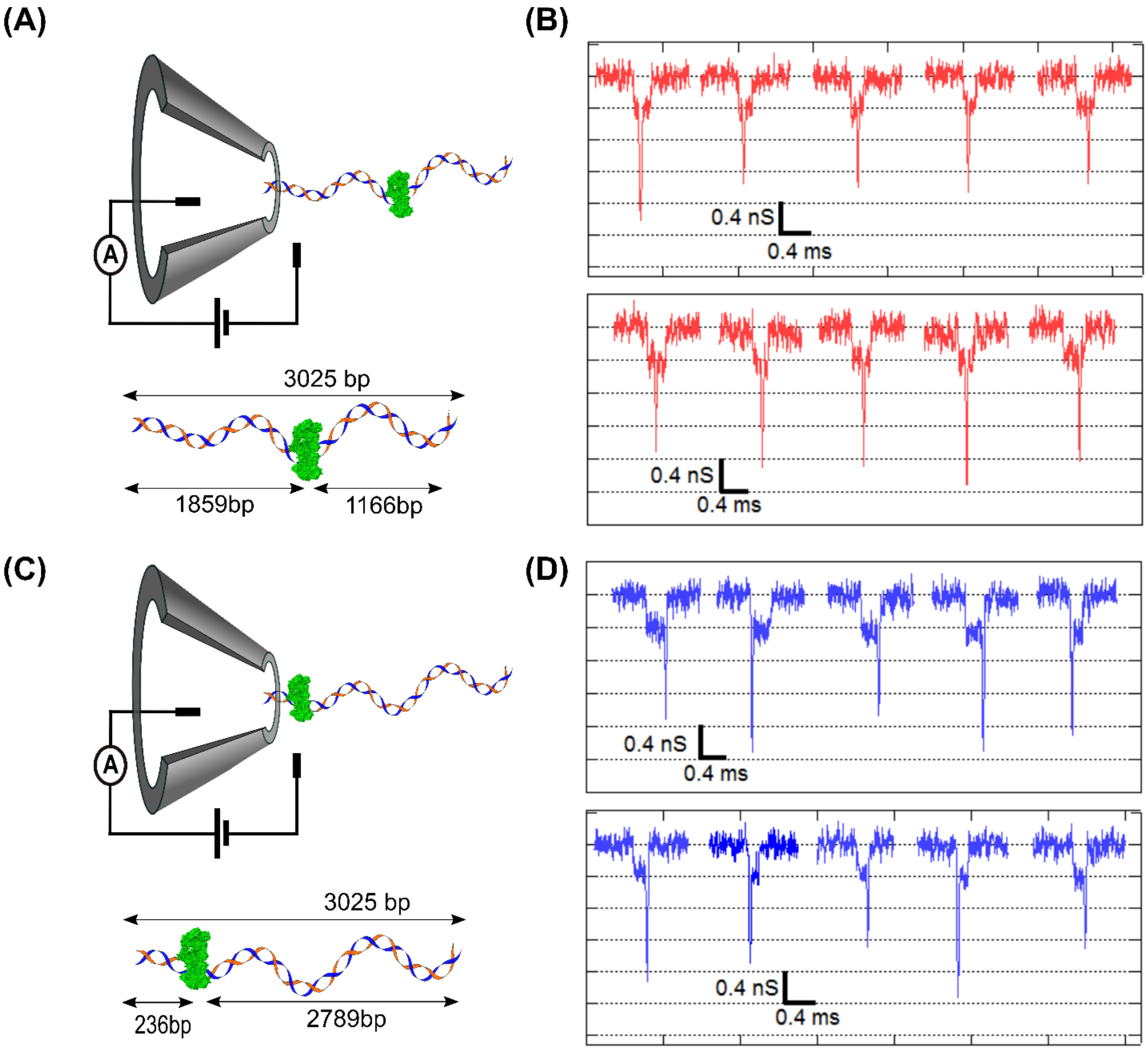


***Figure S8: Distinct signatures of Cas9 bound at different target locations on 3kb DNA.*** *(A) and (B) shows schematics and representative events of DNA- Cas9 complex bound at the center of the target DNA. (C) and (D) shows the same where the DNA- Cas9 complex is bound at the edge of the 3kb DNA.*

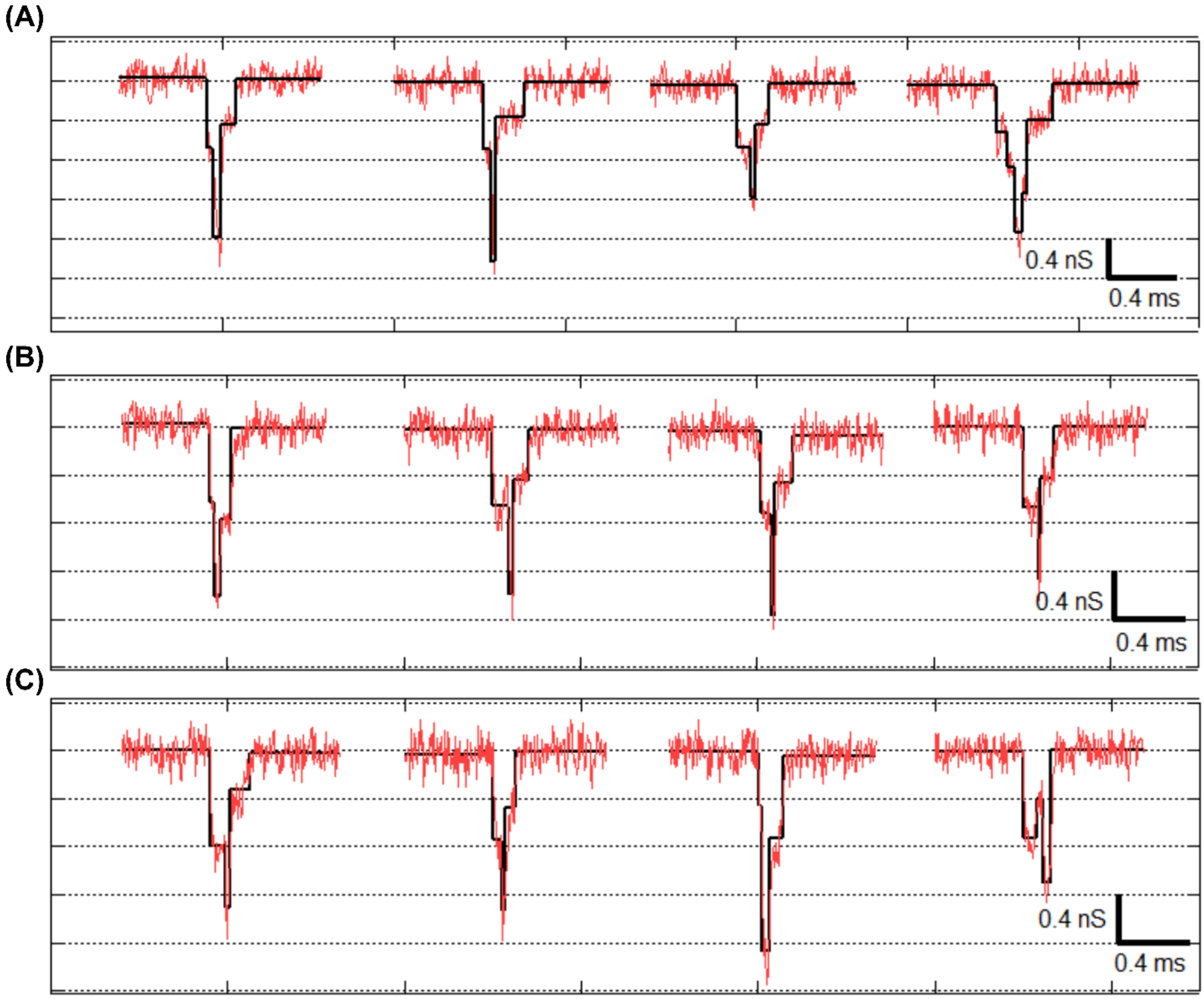


***Figure S9: Representative events showing translocation of folded Cas9-DNA complex(center-target).*** *(A) – (C) shows events with protein spikes during translocation of Cas9 bound to folded DNA. Black line fits show different folding levels of the DNA when Cas9-DNA complex translocate through the pore. Note, no multi-level events (#folds >1) were seen during translocation of control 3 kb DNA (see Figure 2B).*

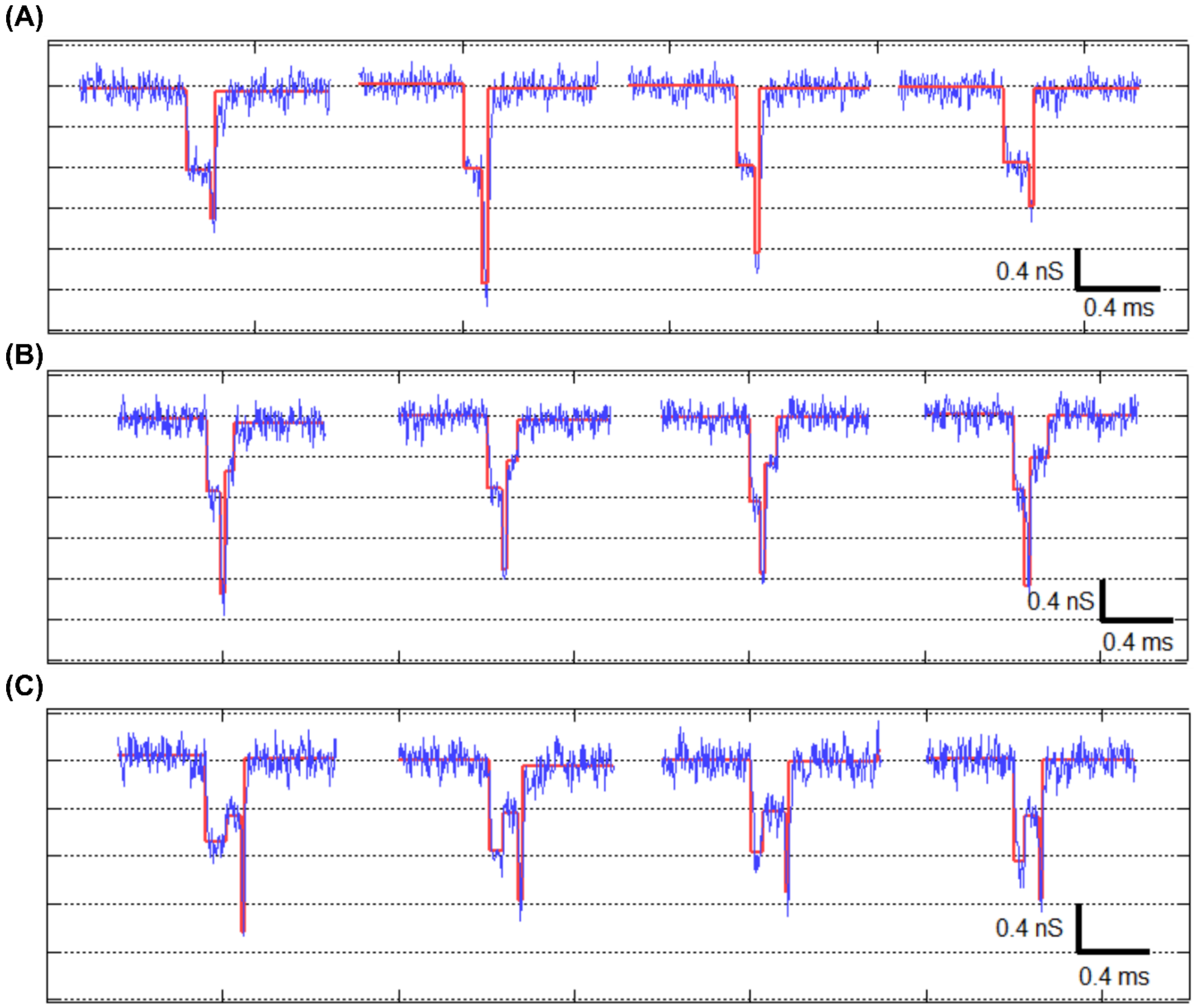


***Figure S10: Representative events showing translocation of folded Cas9-DNA complex (edge-target).*** *(A) – (C) Representative events showing protein spikes during translocation of Cas9 bound folded DNA. Red line fits show different folding levels of the DNA when Cas9-DNA complex translocate through the pore.*

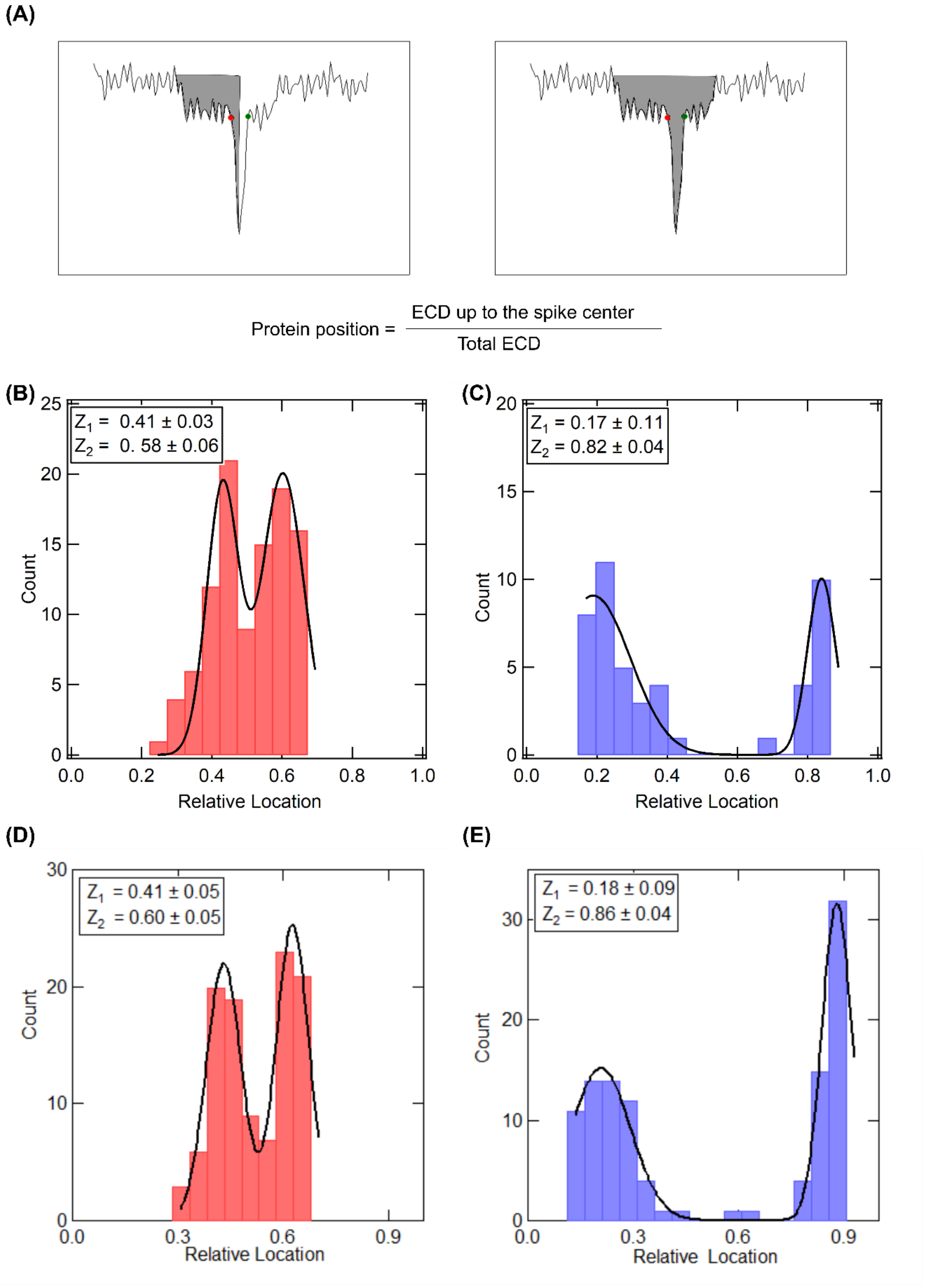


***Figure S11: Location estimation of Cas9 bound on 3025bp DNA.*** *(A) shows in schematic the protein spike (start and end marked with red and green dots respectively) on top of a typical DNA translocation event. Protein position is estimated by comparing ECD of the event up to the spike center (shown as shaded region on the left) to the ECD of the entire event (shaded region on the right). (B)-(C) and (D)-(E) reproduces histograms of Cas9 location on center and edge target DNA, respectively, as measured back-to-back on independent single nanopores.*

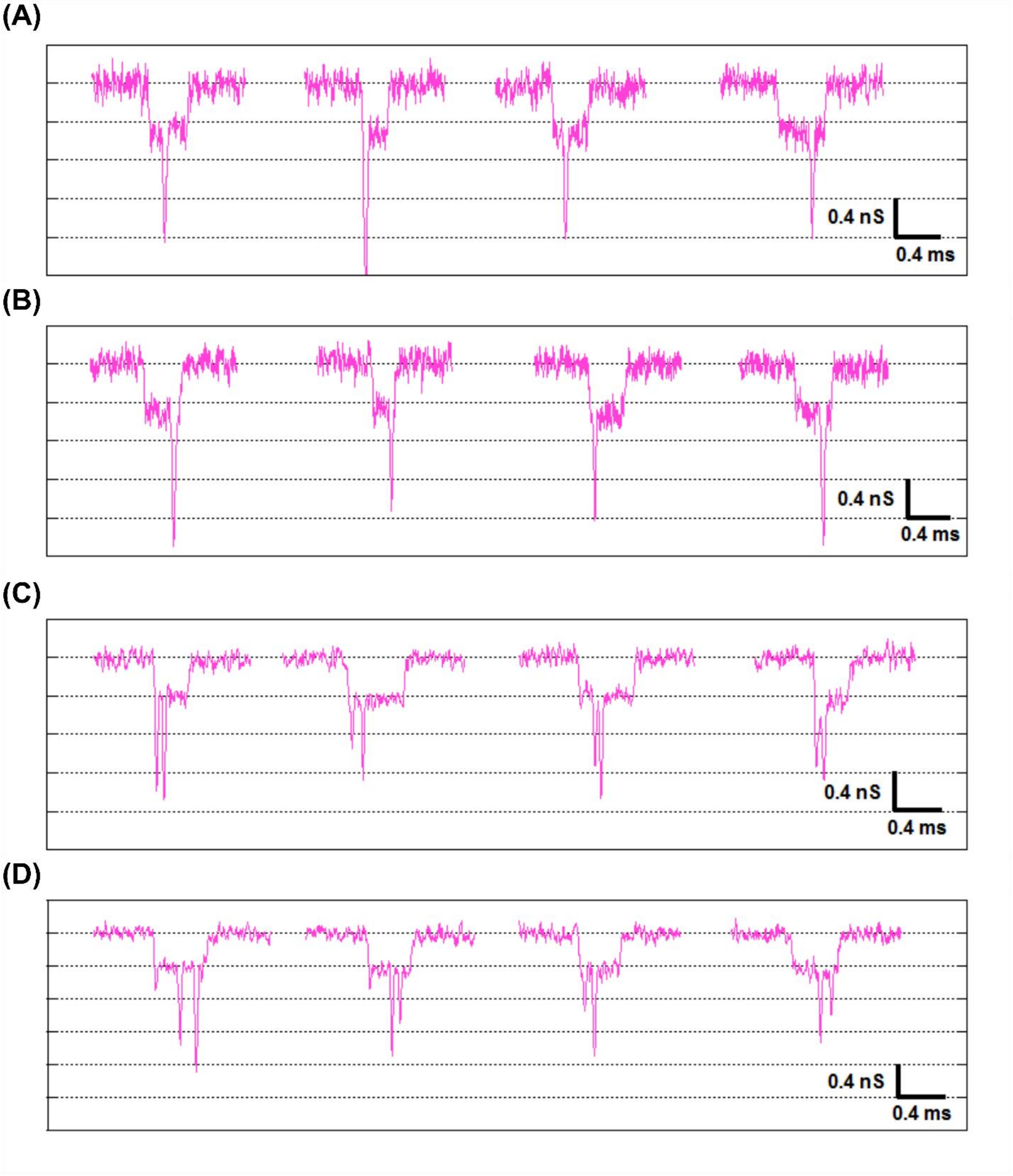


***Figure S12: Representative events of translocation of Cas9-DNA array.*** *(A) & (B) shows translocation of single cas9 bound at multiple location of the 12 available sites. (C) & (D) shows translocation of two cas9 protein bound at multiple location of the 12 available target sites.*

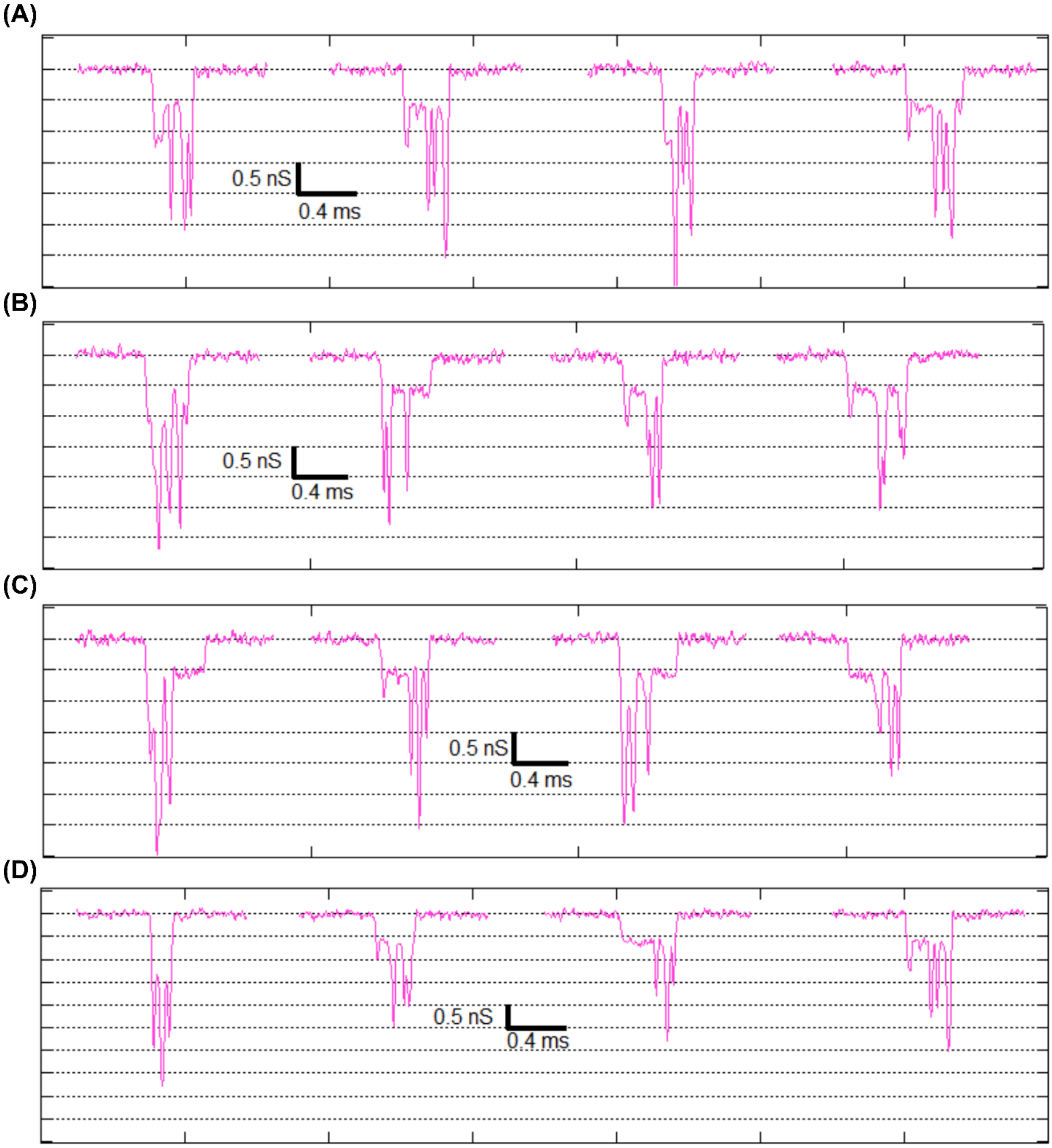


***Figure S13A: Representative events of translocation of Cas9-DNA array.*** *(A), (B), (C) & (D) shows the translocation of at least three cas9 protein bound at different location on 5094 bp DNA having 12 target sites. Note selected events show Cas9 bound to folded and unfolded conformation.*

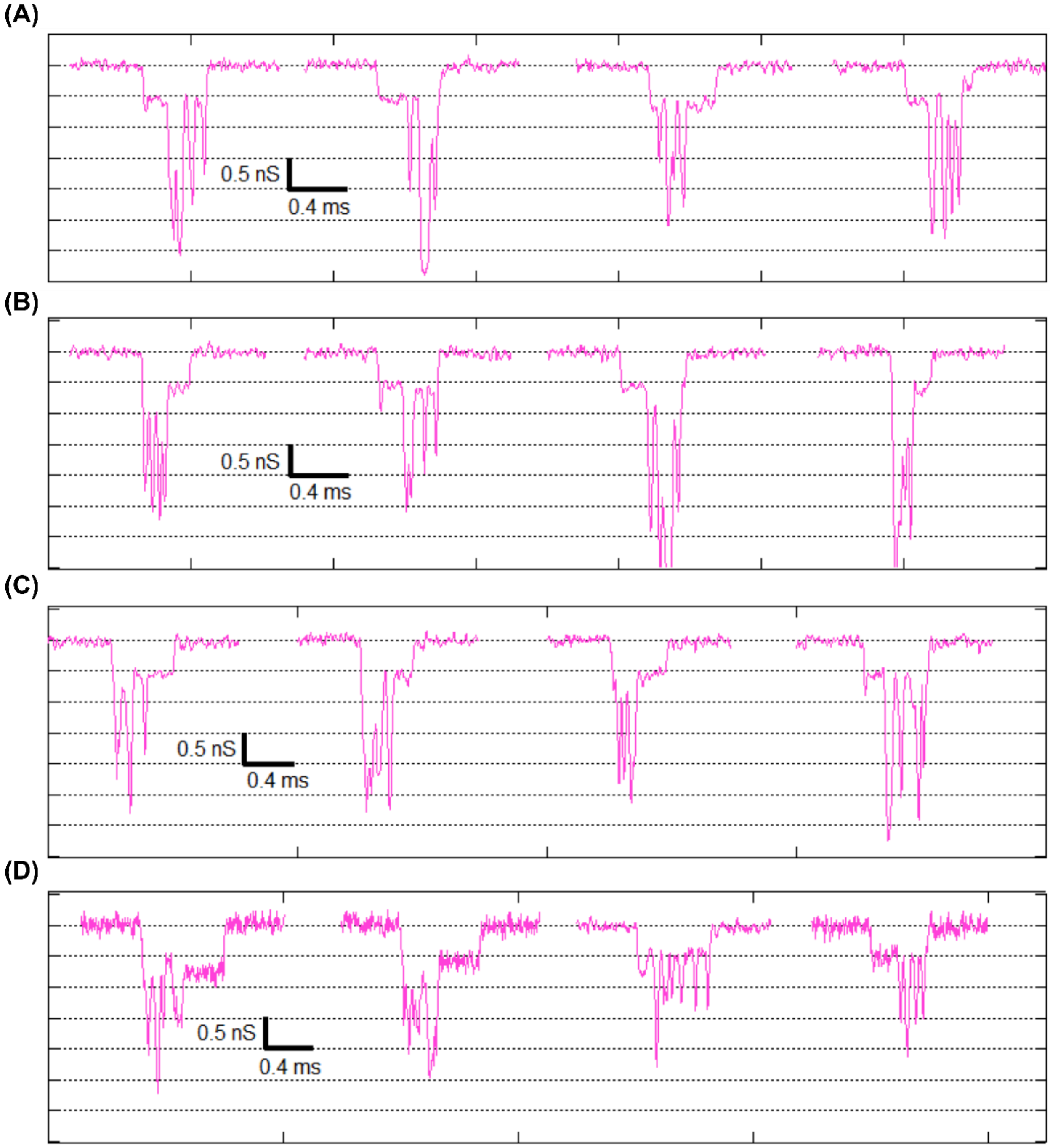


***Figure S13B: Representative events of translocation of Cas9-DNA array.*** *(A), (B), (C) & (D) show the translocation of complex molecules with more than 3 Cas9 protein bound to the 12 possible target sites on the 5094 bp DNA. Note representative events show multi-Cas9 bound in both linear and folded DNA.*

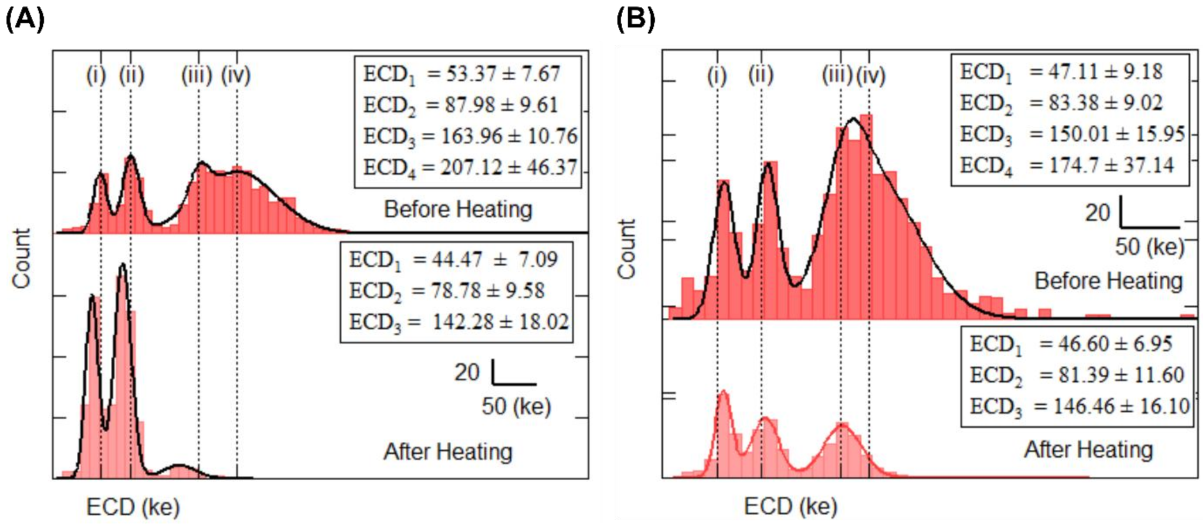


***Figure S14: histogram of ECD for the translocation of DNA Cas9 complex with and without heat treated sample through the nanopore.*** *A (Top) & (bottom) shows ECD histogram for translocation of complex before and after treated with heat performed on pore ID 20. B (Top) & (bottom) shows the reproducible results on pore ID 21.*

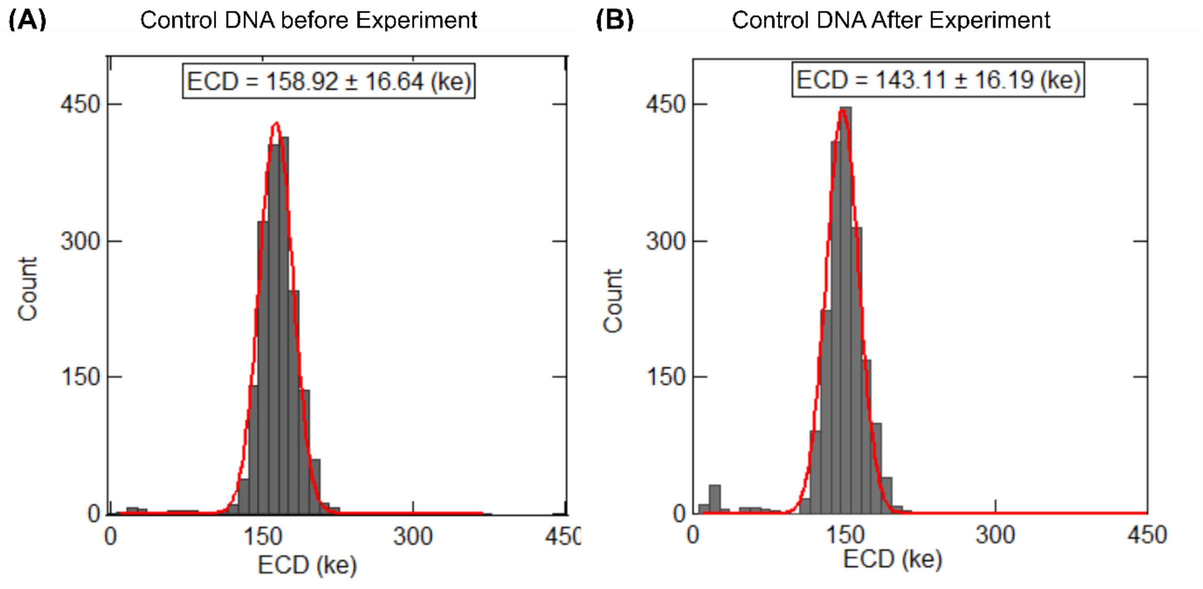


***Figure S15: ECD histogram of control DNA sample.*** *Control DNA sample was measured, in the same nanopore, both before and after the Cas9-DNA complex (bound and heat-released) measurement. Small change in the ECD peak value might be due to pore expansion during the long experiment.*

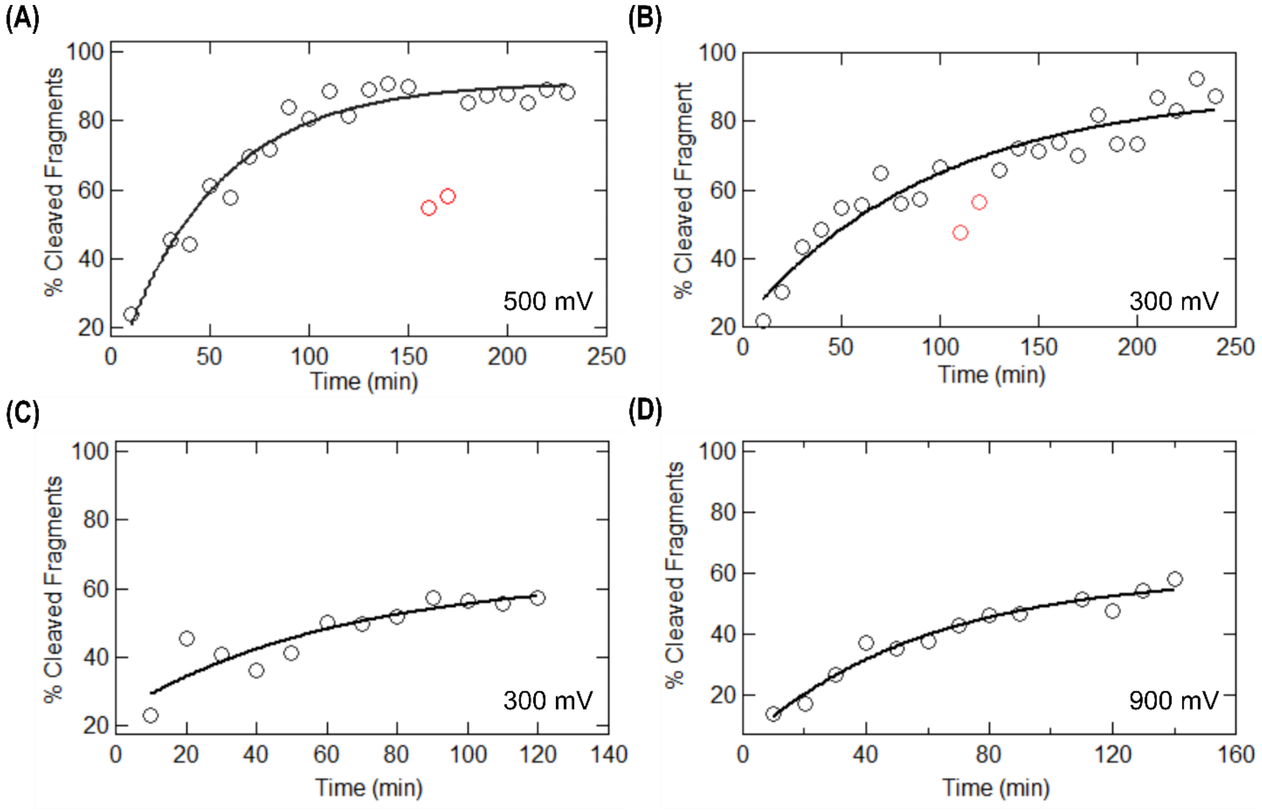


***Figure S16: Rate of release of cleaved DNA fragment in NPB buffer.*** *(A-D) shows the percentage of cleaved fragment release with increasing time in salt, measured in different pores at different voltages. Black curve is first order rate equation fitted to data. During these long experiments, the nanopore can transiently clog affecting the experiment. These time points (shown in red for completeness) were excluded from the fitting.*

First order rate kinetics for release products:

Let us consider the total fraction of complex (C), cleaved fragments (f) and unbound DNA ($D_{free}$), that makes up the reaction mix.

At any time, t

$C + D_{free} + f = 100$

During product release:

$C \rightarrow f$ yields

$-\frac{dC}{dt} = kC$ where k is the rate constant. Solution of above equation is:

$C = C_0 e^{-kt}$

where $C_0$ is complex fraction at t = 0.

This yields our original equation as:

$f = 100 - C_0 e^{-kt} - D_{free}$
